# Evolutionary velocity with protein language models

**DOI:** 10.1101/2021.06.07.447389

**Authors:** Brian L. Hie, Kevin K. Yang, Peter S. Kim

## Abstract

Predicting the order of biological homologs is a fundamental task in evolutionary biology. For protein evolution, this order is often determined by first arranging sequences into a phylogenetic tree, which has limiting assumptions and can suffer from substantial ambiguity. Here, we demonstrate how machine learning algorithms called language models can learn mutational likelihoods that predict the directionality of evolution, thereby enabling phylogenetic analysis that addresses key limitations of existing methods. Our main conceptual advance is to construct a “vector field” of protein evolution through local evolutionary predictions that we refer to as evolutionary velocity (evo-velocity). We show that evo-velocity can successfully predict evolutionary order at vastly different timescales, from viral proteins evolving over years to eukaryotic proteins evolving over geologic eons. Evo-velocity also yields new evolutionary insights, predicting strategies of viral-host immune escape, resolving conflicting theories on the evolution of serpins, and revealing a key role of horizontal gene transfer in the evolution of eukaryotic glycolysis. In doing so, our work suggests that language models can learn sufficient rules of natural protein evolution to enable evolutionary predictability.

## Introduction

Predicting evolutionary order has diverse applications that range from tracing the progression of viral outbreaks to understanding the history of life on earth [1]–[6]. For protein evolution, this prediction is often based on reconstructing and rooting phylogenetic trees of protein sequences [7]. While useful, ordering sequences based on a phylogenetic tree has a number of limiting assumptions; for example, determining the root of the tree can drastically alter the predicted order [8], but beyond the strictest assumptions, determining this root requires manual expertise or external evidence (for example, based on known sampling times or the fossil record), which may not always be available [8], [9].

Here, we propose a novel approach to analyzing and ordering the trajectories of protein evolution that we refer to as “evolutionary velocity,” or “evo-velocity.” Evo-velocity is conceptually inspired by work in theoretical biology that understands evolution as a path that traverses a “fitness landscape” based on locally optimal decisions [2]–[4], [10]–[12]. Our key conceptual advance is that by learning the rules underlying *local* evolution, we can construct a *global* evolutionary “vector field” that we can then use to: (i) predict the root (or potentially multiple roots) of observed evolutionary trajectories, (ii) order protein sequences in evolutionary time, and (iii) identify the mutational strategies that drive these trajectories.

To make local evolutionary predictions, we leverage recent advances in the ability of machine learning algorithms called language models to predict the effects of single-residue mutations on biological fitness when trained on natural sequence variation alone [13]–[16]. Thus far, however, language models have only been applied to modeling local evolution, such as single-residue mutations, rather than more complex changes that occur over long evolutionary trajectories.

Evo-velocity is aimed at closing the gap between landscape-based evolutionary theory [2], [3], [10] and the analysis of evolutionary trajectories observed in nature. Our algorithm is general (we use a single model for all proteins), does not have many of the assumptions typical of phylogenetic methods (for example, evo-velocity can produce multiple roots or model convergent evolution), and requires sequence data alone. We use evo-velocity to analyze protein evolution across a breadth of organisms and evolutionary timescales—from the evolution of viral proteins over the course of years to the evolution of enzymes across all three domains of life— suggesting how we might expand our ability to understand and predict evolution.

## Results

### Overview of language models and evo-velocity

Our approach is based on the premise that evolution occurs through locally optimal changes that preserve or enhance evolutionary fitness, which has theoretical precedent in the concept of a path through a fitness landscape [2], [10]. In theory, predicting local evolution should therefore provide insight into global evolution as well (**Figure 1A**).

**Figure 1:**
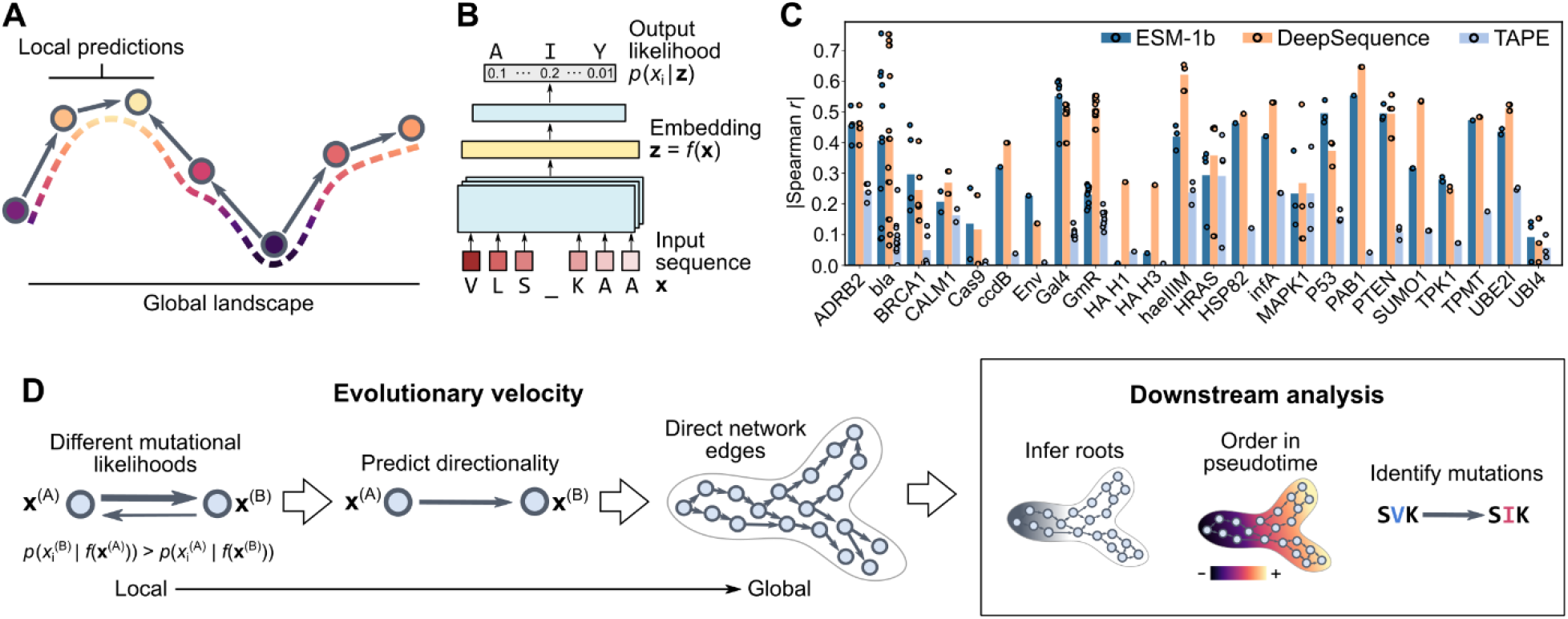
Constructing an evolutionary vector field by predicting local evolution. (**A**) A global evolutionary landscape can be approximated by a composition of local evolutionary predictions. (**B**) To make these predictions, we can leverage language models that learn the likelihood of an amino acid occurring within some sequence context. (**C**) The pseudolikelihoods learned by language models correlate with DMS-based measurements of various notions of protein fitness without the language models being explicitly trained on this data (**Methods**). While DeepSequence trains a separate model for each protein family, ESM-1b and TAPE are general language models each trained on a single, non-redundant dataset. Circles indicate correlations of different DMS profiles within the same study (**Data S1**); bar height indicates the mean across these profiles. (**D**) Evo-velocity uses language model likelihoods to assign a directionality to edges in a sequence similarity network, enabling downstream analysis like predicting root nodes, ordering nodes in pseudotime, and identifying mutations associated with the largest changes in evo-velocity (**Methods**).

To predict the local rules of evolution, we leverage protein language models, which learn the likelihood that a particular amino acid residue appears within a given sequence context (**Figure 1B**). When trained on large corpuses of natural sequences, this language model likelihood is a strong correlate of the effects of mutations on various notions of protein fitness. For example, the ESM-1b language model [15], trained on ~3 million sequences in the UniRef50 database [17] (**Table S1**), can predict the effects of single-residue mutations as quantified by deep mutational scanning (DMS) of diverse proteins [18], [19] (**Figure 1C** and **Data S1**; **Methods**). Surprisingly, this correlation is comparable to that of a state-of-the-art mutational effect predictor [20] that was specially trained on sequence variation within individual protein families (**Figure 1C**); in contrast, ESM-1b is trained on a dataset that removes most intra-family sequence variation [17].

Our key hypothesis is that the likelihoods learned by these large-scale protein language models can be used to provide a notion of directionality within evolutionary trajectories. In our approach, which we call evo-velocity, we first model the “landscape” or the “manifold” [21] of sequence variation by constructing a sequence similarity network [22] in which each node represents a protein sequence and edges connect similar sequences (**Figure 1D**). We quantify sequence similarity as the Euclidean distance in language model embedding space, which can encode complex functional relationships [13]–[16], and we construct the network by connecting a sequence to its *k*-nearest neighbors (KNN), which has been useful in modeling biological landscapes in many genomics applications [23]–[25].

Then, language models assign a directionality to each edge in the KNN network based on the change in language model likelihood between the two sequences in that edge (**Figure 1D**). We hypothesize that evolution moves toward higher likelihoods, which are correlated with higher fitness (**Figure 1C**). Across the entire network, we can then analyze the “flow” of evolution, which includes estimating the root sequences in the network (equivalent to finding the “valleys” of the landscape), ordering sequences in pseudotime (a continuous score that enables rank-based comparison among sequences) [26], visualizing the trajectory in two dimensions [24], and identifying mutations that correlate with the direction of evo-velocity (**Figure 1D**); we provide detailed methodology in **Methods**. Intuitively, the local predictions of language models assign a “velocity” to pairs of sequences that we assemble into an evolutionary “vector field” [27]. In this paper, we implement evo-velocity with a *single* masked language model, ESM-1b, but our framework can readily generalize to other implementations as well.

### Evo-velocity of influenza A nucleoprotein

As initial validation, we used evo-velocity to reconstruct the evolution of the nucleoprotein (NP) of influenza A virus. NP is an excellent evolutionary test case since its sequence evolution is densely sampled through influenza viral surveillance and it undergoes natural selection in the form of host immune pressure, but is less mutable than other viral proteins with a mutation rate of about one amino-acid residue per year [28]. We obtained 3,304 complete NP sequences sampled from human hosts, constructed the sequence similarity network, and computed evo-velocity scores. When we visualized this network in two dimensions [24], we observed phylogenetic structure corresponding to both the sampling year and influenza subtype (**Figures 2A** and **S1A**). Strikingly, the evo-velocity flow through the network (**Methods**) corresponded to the known temporal evolution of NP (**Figure 2A**).

**Figure 2:**
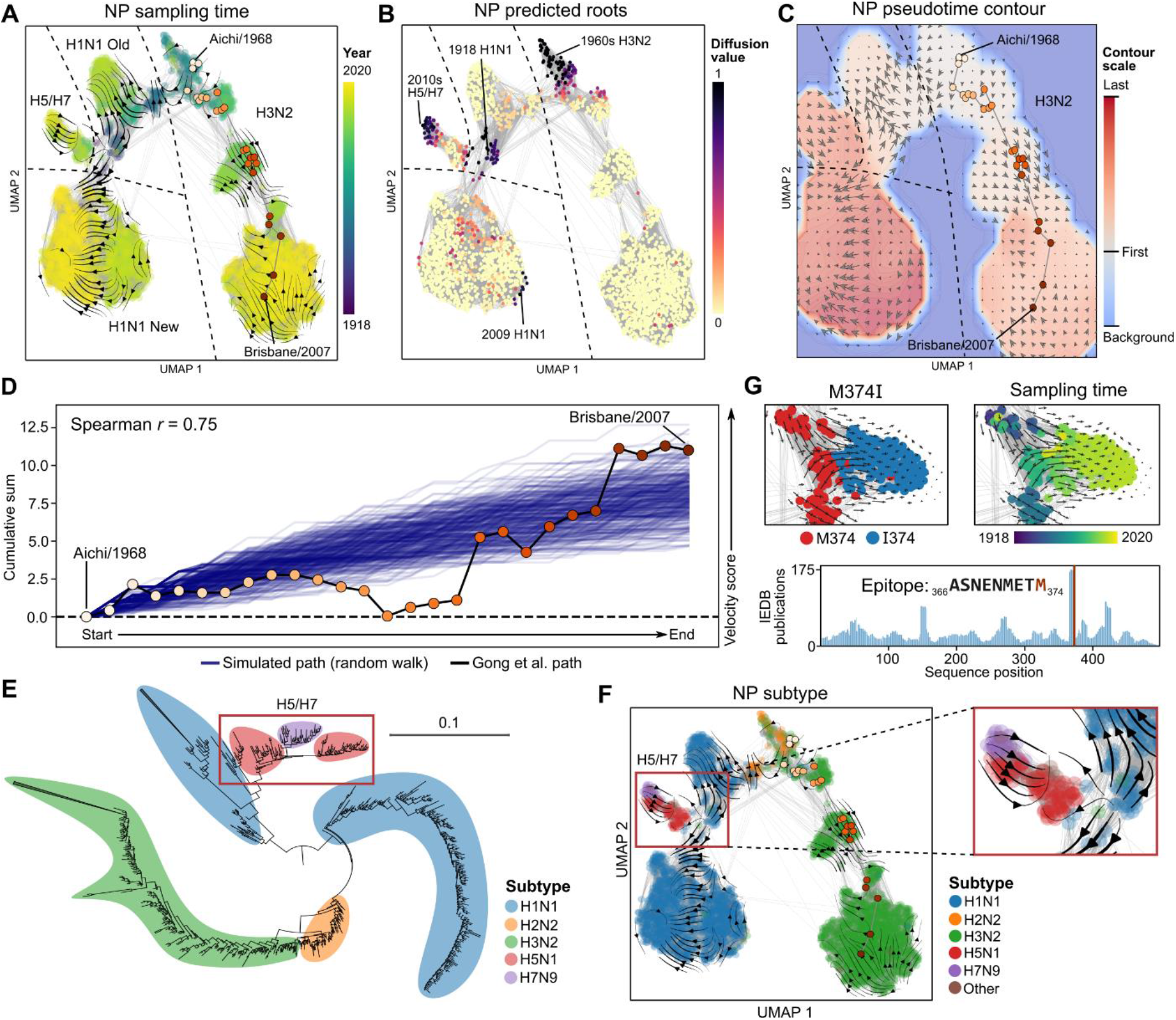
Evo-velocity of influenza A nucleoprotein. (**A**) The landscape of NP sequences, represented as a KNN sequence similarity network, shows structure corresponding to temporal evolution of various subtypes of influenza (**Figure S1A**); gray lines indicate network edges. Overlaying evo-velocity on the visualization as a streamplot shows a visual correlation between the flow of evo-velocity and known sampling time. A known phylogenetic path (orange circles) from Gong et al. [28] starting with Aichi/1968 and ending with Brisbane/2007 moves in the direction of evo-velocity. (**B**) Using the evo-velocity directionality to predict roots reveals four main root regions corresponding to the beginnings of different influenza pandemic events throughout history. (**C**) Ordering sequences in pseudotime and visualizing pseudotime values in a two-dimensional contour plot shows pseudotime increase in the direction of evo-velocity, which here is visualized as a two-dimensional field of evo-velocity vectors. (**D**) On average, the Gong et al. path visualized in (**A**) and (**C**) has positive changes in evo-velocity scores over time and largely resembles simulated paths generated by performing random walks across our evo-velocity landscape (**Methods**). A portion of the Gong et al. path with negative evo-velocity scores may be due to ordering ambiguities that are better resolved by considering evo-velocity. (**E**) A maximum-likelihood, midpoint-rooted phylogenetic tree of all NP sequences conveys that H5N1 and H7N9 subtype sequences branch off from H1N1 sequences. (**F**) In contrast, evo-velocity predicts an independent origin of H5N1/H7N9 influenza [29] (see **B**) and sequence similarity with H1N1 due to convergent evolution. (**G**) The M374I mutation to NP has the second strongest magnitude change in evo-velocity (**Methods**) and is located in the most well-studied human T-cell epitope on NP (**Table S2**).

Since visualizing this flow in two dimensions can be prone to information loss or distortion through dimensionality reduction [27], we sought to further quantify the relationship between evo-velocity and NP evolution. We first verified that, on average, the evo-velocity scores of the individual network edges increase along with greater differences in sampling time (**Figure S1B**). We then quantified global evo-velocity patterns using a diffusion analysis to estimate the network’s roots (**Methods**). Interestingly, the evo-velocity-inferred root sequences corresponded to the main species-crossover events in influenza history (**Figure 2B**), suggesting that our analysis accurately inferred the evolutionary origins of NP as observed in human hosts. We then used these roots to order sequences according to evo-velocity pseudotime (**Methods**) and observed a significant correlation between pseudotime and known sampling date (Spearman *r* = 0.49, two-sided *t*-distribution *P* = 4 × 10^−197^) (**Figure 2C**). We also observed that a well-characterized phylogenetic path of NP [28] progressed, on average, in the same direction as the evo-velocity gradient (**Figure 2A,C**) and agreed with simulated paths generated by random walks across our evo-velocity landscape (**Figure 2D**; **Methods**).

When comparing our evo-velocity landscape to a standard phylogenetic tree, we observed that evo-velocity can model more complex evolutionary relationships. For example, a midpoint-rooted phylogenetic tree of all NP sequences (**Methods**) visually suggests that the H5N1- and H7N9-subtype sequences branch off from H1N1 (**Figure 2E**). Instead, evo-velocity predicts an independent origin of H5N1/H7N9 (**Figure 2C,F**), consistent with epidemiological data indicating recent zoonotic crossover of H5 and H7 avian influenza [29]. Evo-velocity also predicts that the observed similarity of H5N1/H7N9 and H1N1 NP sequences sampled in human hosts is due to convergent evolution (**Figure 2F**), which is challenging to explicitly represent with a phylogenetic tree.

We next sought to use our evo-velocity landscape to provide new insight into NP evolution. We therefore identified the mutations that corresponded to the strongest changes in the evo-velocity scores (**Methods**). Of the top five such mutations in NP, all are present in experimentally-validated human T-cell epitopes and one of these mutations, M374I, is located in the most well-characterized linear NP epitope in the Immune Epitope Research Database (IEDB) [30] (**Figures 2G**, **S1C**, and **Table S2**). Moreover, all five mutations involve a single-nucleotide substitution resulting in a methionine changed to a hydrophobic or polar-uncharged amino acid residue, suggesting a possible T-cell escape strategy that has recurred in multiple NP epitopes throughout history (**Figures 2G** and **S1C**).

All NP sequences in our analysis belong to a single UniRef50 sequence cluster [17] for which the representative sequence is from a 1934 H1N1 virus (**Figure S1D**). We found that similarity to sequences present in UniRef50, the ESM-1b training dataset, does not explain evo-velocity pseudotime (**Table S3**; **Methods**). We also found that computing evo-velocity scores with a smaller language model, TAPE [14], trained with a different model architecture on the Pfam database of protein families [31], closely reproduced the ESM-1b evo-velocity results (Spearman *r* = 0.93, two-sided *t*-distribution *P* < 1 × 10^−308^) (**Table S4** and **Figure S1E,F**). Using simpler evolutionary scores to compute velocities or using binary sequence embeddings also largely reproduced the ESM-1b results, though with weaker temporal correlation (**Figure S1G** and **Table S5**; **Methods**). Together, these results suggest that our evo-velocity results are not explained by trivial language model preference to UniRef50. We also found that evo-velocity pseudotime was not explained by variation in sequence length (**Table S6**).

Evo-velocity was therefore able to reconstruct the direction of NP evolution without any explicit knowledge of influenza subtype or when the NP sequences were sampled. Moreover, we found that the generic rules learned by large language models were sufficient to predict the evolution of a specific protein.

### Evo-velocity of viral proteins

Given the promising results for NP, we were therefore interested in seeing if evo-velocity could generalize to other viral proteins as well. We next analyzed the evolution of influenza A hemagglutinin (HA), a more variable protein on the viral surface responsible for viral-host membrane fusion [32]. As with NP, evo-velocity analysis of 8,115 HA sequences recovered roots corresponding to the known origins of HA H1 in humans from 1918 and 2009 H1N1 pandemics, and evo-velocity pseudotime was strongly correlated with sampling date (Spearman *r* = 0.63, two-sided *t*-distribution *P* < 1 × 10^−308^) (**Figure 3A,B**). Despite the higher sequence variability of HA than NP, evo-velocity was still able to reconstruct the trajectory and directionality of HA evolution.

**Figure 3:**
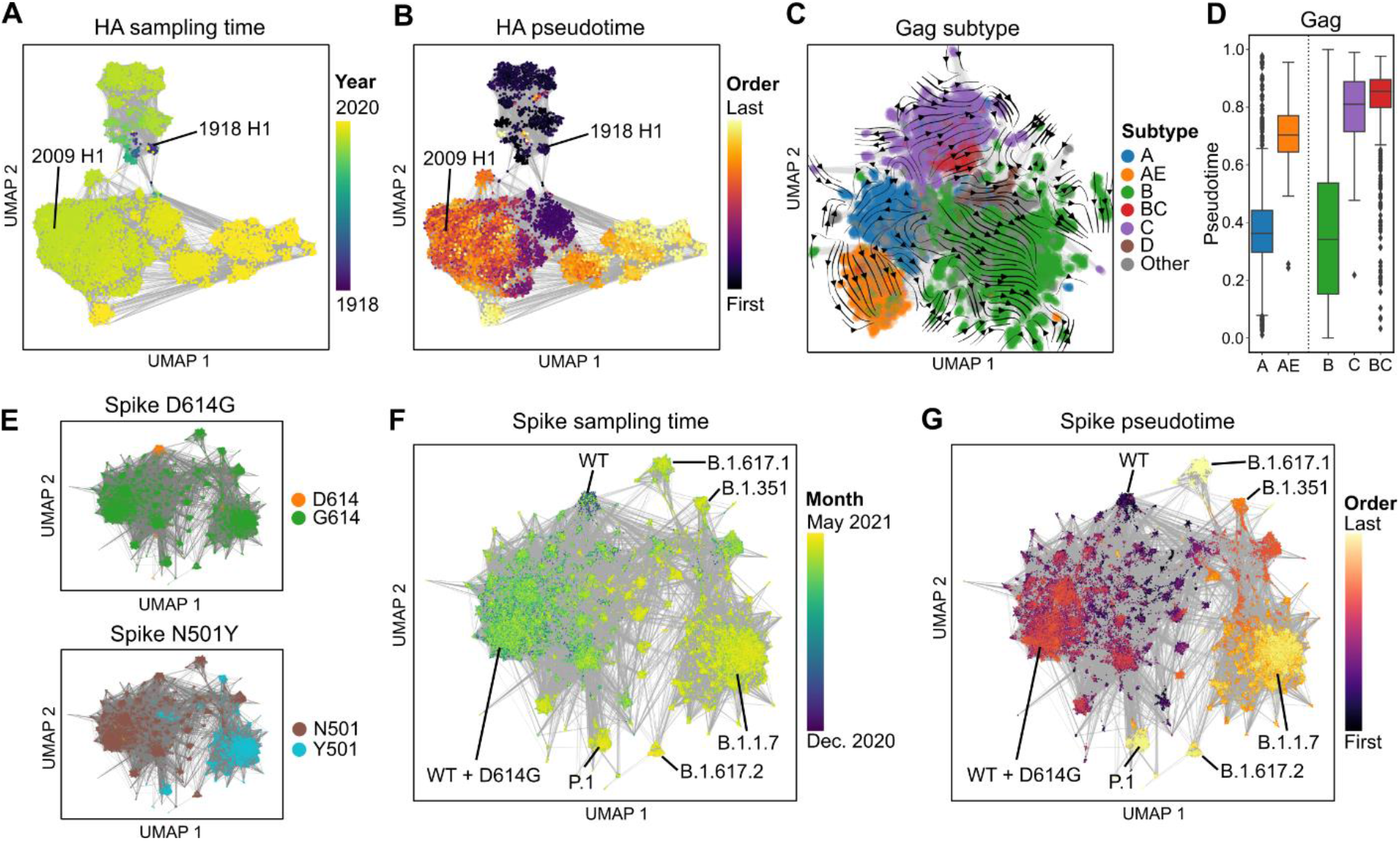
Evo-velocity of viral proteins. (**A, B**) Temporal evolution of HA H1 evolution is captured in the UMAP landscape and is also predicted by evo-velocity pseudotime. Two main clusters correspond to the two main pandemic trajectories of H1N1, the first beginning in the early twentieth century and the second beginning in the early twenty-first century. (**C, D**) An evo-velocity streamplot of Gag evolution illustrates the branching trajectories of HIV-1 subtypes, including major subtypes like A, B, and C preceding circulating recombinant forms like AE and BC. Box extends from first to third quartile with line at the median, whiskers extend to 1.5 times the interquartile range, and diamonds indicate outlier points. (**E-G**) Variants of Spike (identified using characteristic mutations like D614G and N501Y) that emerge in later portions of the COVID-19 pandemic are also predicted to be later in evo-velocity pseudotime.

As with NP, our HA pseudotime results were not explained by sequence similarity to the training dataset (**Figure S2A** and **Table S3**). We were also able to use TAPE-based velocities to identify similar root regions in the post-2009 pandemic trajectory, but TAPE had a more difficult time identifying the 1918 sequences as oldest, most likely due to TAPE’s smaller model size and less capable mutational effect predictions (**Figures 1C**, **S2B-D**, and **Table S4**).

We next analyzed the evolution of the group specific antigen (Gag) polyprotein of human immunodeficiency virus type 1 (HIV-1) using 18,018 sequences. Visualizing the sequence similarity network overlaid with evo-velocity reveals a flow corresponding to the known subtype branching history of HIV-1, with circulating recombinant forms (for example, subtypes AE and BC) branching off of the main subtypes and occurring later in pseudotime (**Figure 3C,D**). HIV-1 Gag sequences also had strong positive velocities compared to phylogenetically-similar Gag sequences from chimpanzee simian immunodeficiency virus (SIVcpz) (**Figure S2E**), consistent with a SIVcpz origin preceding the evolution of pandemic HIV-1 [33]. We observed much weaker correlation between pseudotime and sampling date (Spearman *r* = 0.093, two-sided *t*-distribution *P* = 5 × 10^−32^) (**Figure S2F**) compared to influenza proteins, consistent with the much weaker population-level immune pressure on Gag evolution. Gag pseudotime was not explained by sequence similarity to UniRef50 (**Table S3**) and was also reproducible using TAPE-based velocities (**Figure S2G** and **Table S4**).

We next applied our algorithm to analyze 46,986 sequences of the Spike glycoprotein of severe acute respiratory syndrome coronavirus 2 (SARS-CoV-2) across a much shorter historical timescale of around eighteen months. The sequence similarity network reconstructs the overall trajectory of Spike evolution, and evo-velocity analysis identifies the sequence clusters associated with later sequences, including the B.1.1.7, B.1.351, B.1.617.1, B.1.617.2, and P.1 variants-of-concern [34], as later in pseudotime (**Figures 3E-G**). Despite a shorter evolutionary timescale, evo-velocity pseudotime and sampling date still had a Spearman correlation of 0.41 (two-sided *t*-distribution *P* < 1 × 10^−308^). We also note that SARS-CoV-2 Spike evolution occurred outside of the temporal range associated with both language model training datasets and we were also able to reproduce the results with TAPE-based evo-velocity (**Figure S2H** and **Table S4**).

Across these four viral proteins, therefore, evo-velocity was able to reconstruct the directionality of evolution consistent with known trajectories. Importantly, all of our analysis was based on a single model that was trained without explicit knowledge of viral sampling date, subtype, or protein-specific sequence variation.

### Evo-velocity of eukaryotic proteins

After validating our approach with known viral trajectories, we wanted to see if evo-velocity could generalize to longer trajectories, such as protein evolution that spans multiple species. Though we have access only to extant sequences, we hypothesized that evo-velocity might still provide useful orderings if some extant sequences are closer to the ancestral sequence than others. As an initial test case, we analyzed the globin protein family due to its extensive phylogenetic characterization [35], including laboratory reconstruction of ancestral intermediates, that we can use to validate our model (**Figure 4A**).

**Figure 4:**
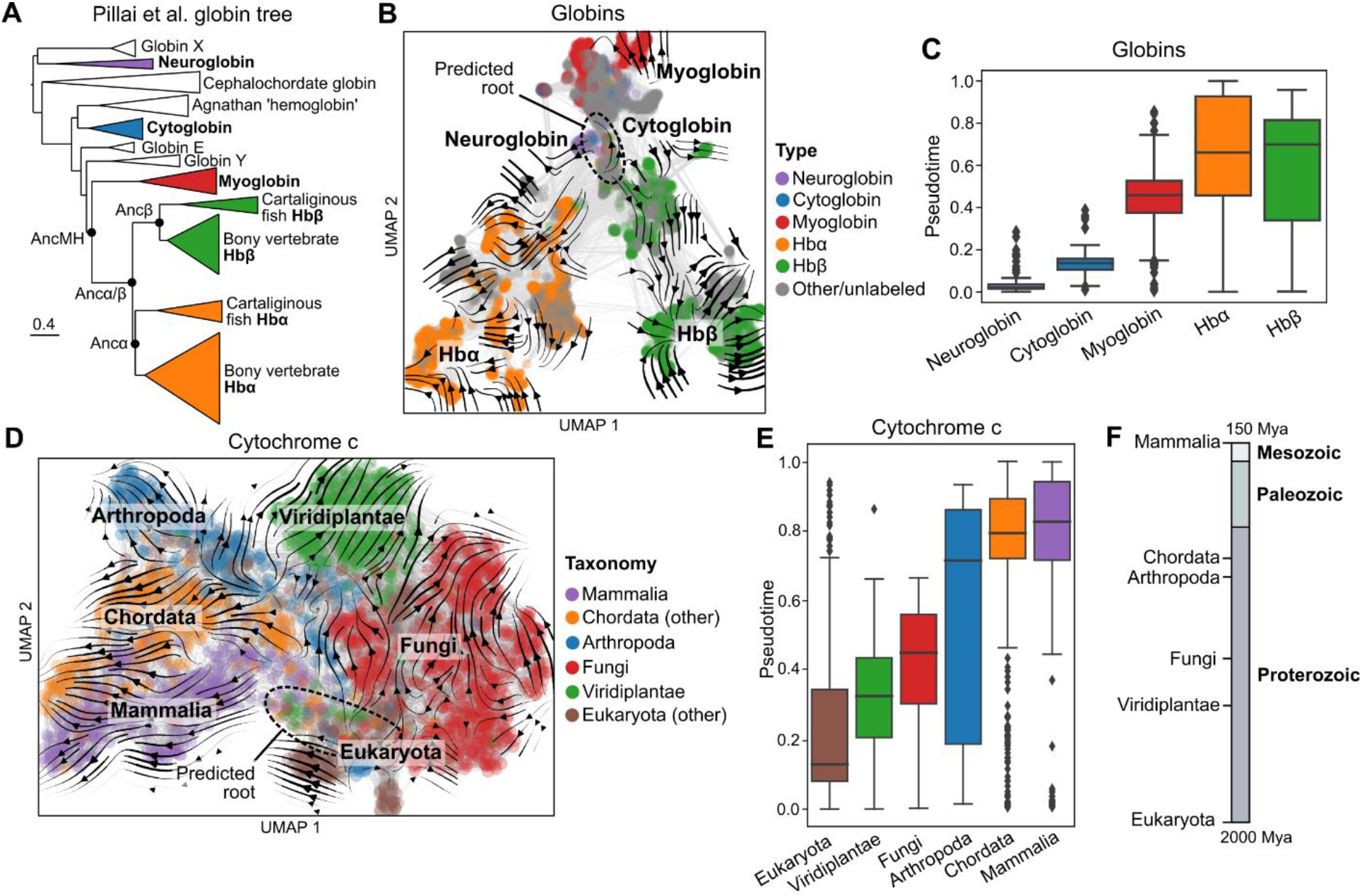
Evo-velocity of eukaryotic proteins. (**A**) The maximum likelihood phylogenetic tree determined by Pillai et al. [35] is rooted in globin X and neuroglobin with the longest branches extending to Hbα and Hbβ. (**B**) The landscape of globin sequences shows a branching trajectory with the predicted root also closest to neuroglobin (**Figure S3A**). (**C**) Computing pseudotime from this predicted root places Hbα and Hbβ as most recent in evolution, consistent with the tree of Pillai et al. (**D**) The landscape of cytochrome c sequences shows clustering structure corresponding to known taxonomic labels, with the evo-velocity gradient beginning among single-celled eukaryotes and plants (**Figure S4A**). (**E, F**) The ordering of the median evo-velocity pseudotimes of various taxonomic labels corresponds to the evolutionary orderings in geologic time determined by molecular clocks and the fossil record [37]. For all boxplots: box extends from first to third quartile with line at the median, whiskers extend to 1.5 times the interquartile range, and diamonds indicate outlier points.

The landscape of 6,097 eukaryotic globin sequences forms a branching trajectory with three major divisions corresponding to myoglobin, alpha hemoglobin, and beta hemoglobin (**Figure 4B**). The predicted root region lies in the part of the landscape closest to neuroglobin and cytoglobin (**Figure S3A,B**). Of the major classes of globins, neuroglobin is estimated to be earliest in pseudotime while the alpha (Hbα) and beta (Hbβ) subunits of hemoglobin occur last in pseudotime (**Figure 4C**), consistent with a previous analysis of globin phylogeny by Pillai et al. [35] (**Figure 4A**). These results are also reproducible when using TAPE to compute the evo-velocity scores (**Figure S3C,D** and **Table S4**) and when controlling for sequence similarity to the training dataset (**Figure S3D** and **Table S3**; **Methods**).

Previous work [35] has also reconstructed ancestral globins that are confirmed to be viable oxygen binders and that progress from a monomeric myoglobin/hemoglobin ancestor (AncMH) to a dimeric alpha/beta hemoglobin ancestor (Ancα/β) to a tetramer formed by separate alpha and beta hemoglobin ancestors (Ancα and Ancβ, respectively) (**Figure 4A**). Consistent with evo-velocity increasing over evolutionary time, the ESM-1b language model likelihood, on average, increases from AncMH to extant myoglobin and hemoglobin sequences, but this improvement diminishes for more proximal ancestors (**Figure S3E**). Together, our globin results suggest that evo-velocity pseudotime within a protein family can recover ordering relationships over longer evolutionary timescales.

To further test this hypothesis, we analyzed 2,128 sequences of cytochrome c, a well-studied protein in evolutionary biology due to its high sequence conservation among most eukaryotes [36]. When visualized, the sequence similarity network combined with evo-velocity reflects the taxonomic diversification of the eukaryota (**Figure 4D**). The ordering of the median pseudotimes of different taxonomic classes also recapitulates their known ordering in geologic time based on estimates from the fossil record and molecular clocks [37] (**Figures 4E,F** and **S4A,B**), and the variation in pseudotime enables a notion of uncertainty in the form of pseudotemporal confidence intervals. We were also able to reproduce pseudotemporal orderings when using TAPE to compute the evo-velocity scores (**Figure S4C,D** and **Table S4**) and when controlling for sequence similarity to the training dataset (**Figure S4D** and **Table S3**). In total, therefore, our analysis of well-studied eukaryotic protein families demonstrates that evo-velocity can generalize to protein evolution at much longer timescales.

### Evo-velocity of ancient evolution

After validating that evo-velocity could reconstruct longer trajectories of protein evolution, we applied evo-velocity to highly-conserved proteins, which often have substantial evolutionary uncertainty [6], to yield new insight into ancient evolution. A protein family with considerable evolutionary uncertainty is that of the serine protease inhibitors, or serpins [38], [39]. Unlike most highly-conserved families, in which most of the diversity is bacterial, most of the diversity among serpins is eukaryotic, which we likewise observe in our landscape of 22,737 serpin sequences (**Figure 5A,B**). This has led to conflicting theories as to whether serpins indeed have a phylogenetic root in eukaryotes, with prokaryotes acquiring serpins via horizontal gene transfer (HGT), or if this root is an artifact of greater eukaryotic diversity biasing phylogenetic root estimation [38]–[40]. Since evo-velocity is not prone to the same bias when estimating roots, we used evo-velocity to order serpin sequences in pseudotime and found that the main predicted root region was located among the prokaryotes (**Figures 5B,C** and **S5A**). These results, along with the uncertain mechanism of eukaryotic-to-prokaryotic HGT [40], provide strong evidence that serpin evolution follows a more canonical trajectory.

**Figure 5:**
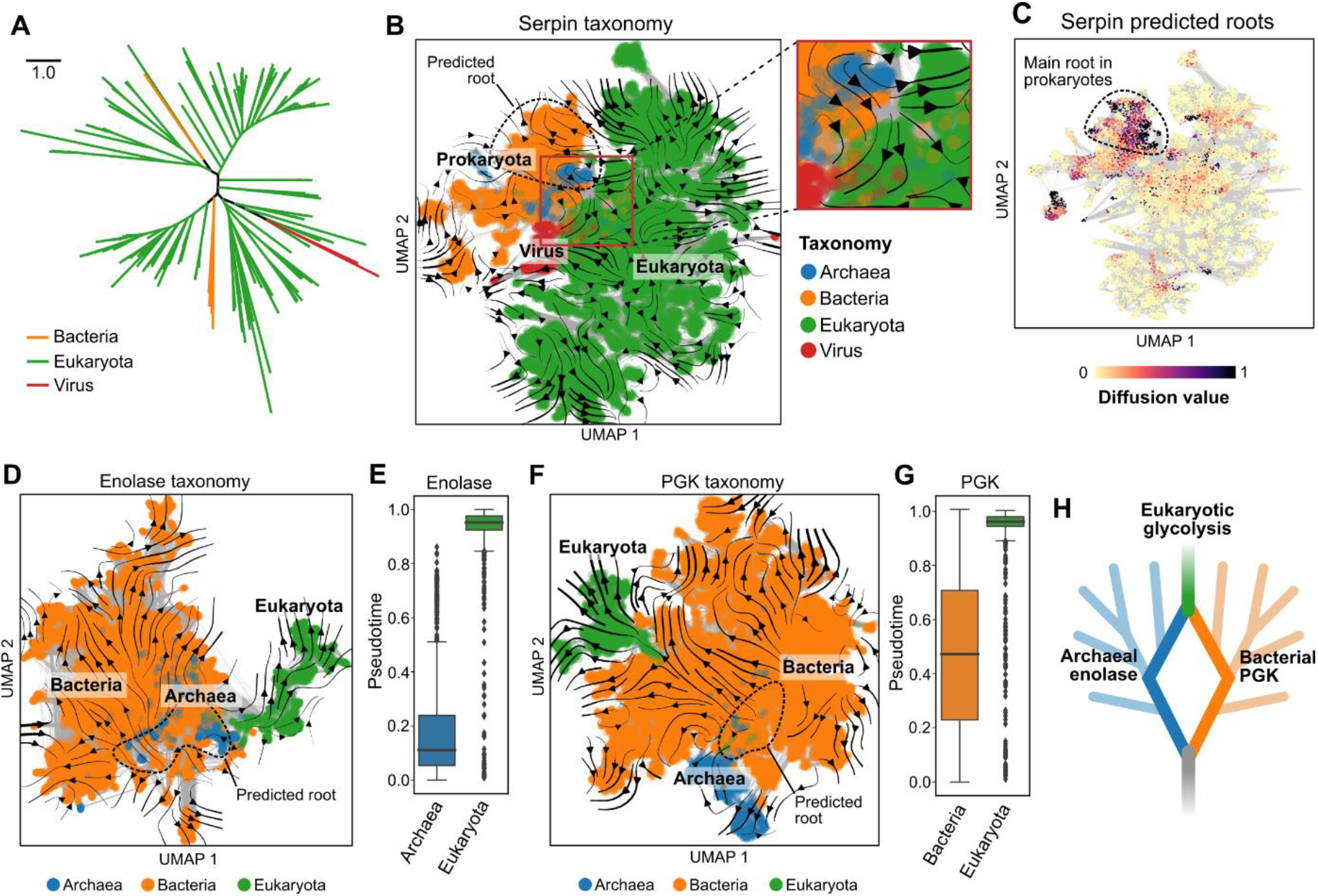
Evo-velocity of ancient evolution. (**A**) The unrooted maximum likelihood phylogenetic tree of serpins shows substantially more eukaryotic than prokaryotic diversity, leading some to hypothesize a eukaryotic root [38], [39]. (**B, C**) Despite lower prokaryotic diversity, evo-velocity still identifies the root of serpins within the prokaryotes, and eukaryotes are the last domain in evo-velocity pseudotime (**Figure S5A**), suggesting that prokaryotic serpins were not acquired from eukaryotes via HGT [38], [39]. (**D, E**) The evo-velocity-predicted root of the enolase landscape begins in a region of archaea and some bacteria, with eukaryotic enolase as the most recent in pseudotime and directly proximal to archaeal enolase on the sequence landscape (**Figure S5B,C,F**). (**F, G**) The evo-velocity-predicted root of the PGK landscape begins in a mostly bacterial region with some archaea, with eukaryotic PGK also very recent in pseudotime and directly proximal to bacterial PGK (**Figure S5D,E,G**). (**H**) The sequence landscapes and evo-velocity-predicted roots suggest that the component enzymes of eukaryotic glycolysis were acquired through different evolutionary paths via HGT; figure adapted from Figure 1 of Weiss et al. [6]. For all boxplots: box extends from first to third quartile with line at the median, whiskers extend to 1.5 times the interquartile range, and diamonds indicate outlier points.

We next analyzed two of the most conserved glycolytic enzymes, enolase and phosphoglycerate kinase (PGK) [41]–[43]. The landscape of 31,901 sequences from the enolase family shows a clear evo-velocity-predicted root region located in bacterial and archaeal sequences (**Figures 5D** and **S5B,C**). Archaea are also oldest in pseudotime and eukaryota are newest, with bacteria showing considerable pseudotemporal variation (**Figures 5E** and **S5C**). The landscape of 30,455 PGK sequences has a similar origin in a region with bacterial and archaeal sequences (**Figures 5F** and **S5D,E**), though with more pseudotemporal variation among archaeal PGK (**Figures 5G** and **S5E**).

The largest difference between the enolase and PGK landscapes lies in the location of eukaryota: while both estimate eukaryota to be recent in pseudotime, eukaryotic sequences branch off of archaeal enolase but branch off of bacterial PGK (**Figures 5D,F**); similar patterns are also observed when visualizing the unrooted phylogenetic trees of both proteins (**Figures S5F,G**). These results suggest an archaeal origin of eukaryotic enolase and a bacterial origin of eukaryotic PGK (**Figure 5H**) and are consistent with HGT contributing to a mixture of archaeal and bacterial genes in the last eukaryotic common ancestor [6]. These results are also consistent with a component-wise evolution of glycolysis [41], rather than the pathway being inherited in totality from a single organism.

In all three highly conserved proteins that we tested, we were able to reproduce evo-velocity pseudotime even when explicitly controlling for sequence similarity to the training dataset (**Figure S5A,H** and **Table S3**) and when using TAPE to compute the evo-velocity scores (**Figure S5A,H** and **Table S4**). Variability in sequence length did not explain evo-velocity pseudotime (**Table S6**). Moreover, the direction of the evo-velocity gradient is not explained by trivial training set bias toward eukaryotes, as most of the sequences in UniRef50 are bacterial (**Table S1**), and we emphasize that no explicit taxonomic information was provided to our algorithm. Rather, our results suggest that evo-velocity can provide insight into evolution at the longest evolutionary timescales.

## Discussion

The degree to which evolution is predictable has been a longstanding debate [3], [4], [44], [45]. Here we show that large-scale protein language models can learn evolutionary rules well enough to predict the ordering of sequences in evolutionary time. While the phylogenetic tree is the oldest conceptual model of evolution [1] and has had wide application to natural sequence variation [7], we show that landscape-based theory [3], [10]–[12] combined with modern algorithms can also provide novel evolutionary insight that is complementary to existing approaches.

Evo-velocity has a number of distinctives with respect to phylogenetic tree reconstruction. Evo-velocity is especially suitable for analyzing large (~1000 or more) collections of sequences. We currently limit our analysis to extant sequences, rather than artificially reconstructing ancestral sequences, though these could be incorporated into the analysis as well. In viewing evolution as a landscape, evo-velocity admits multiple “valleys” that we refer to as roots. Because we predict the directionality of edges in the network, evo-velocity roots are also better mathematically determined than phylogenetic roots [9], [46] (though users could manually specify root sequences as well). Evo-velocity landscapes can also better model phenomena like convergent evolution (**Figure 2F**).

We also find that evo-velocity provides a helpful notion of uncertainty in its predictions that is less natural to obtain from standard phylogenetic methods. For example, evo-velocity reports multiple roots, indicating evolutionary ambiguity regarding the oldest sequences or reflecting discontinuous trajectories due to missing evolutionary ancestors. Similarly, the most robust ordering relationships are at the level of groups of sequences, providing pseudotemporal confidence intervals.

Computationally, our results are striking in that a single language model trained on diverse, natural protein sequences seems to learn generic evolutionary rules. This is corroborated by our finding that two independently-trained language models, ESM-1b and TAPE, can produce very similar pseudotemporal ordering results (**Table S4**), even though TAPE is a much weaker mutational effect predictor than ESM-1b (**Figure 1C**). The robustness of evo-velocity pseudotime to language model implementation may be because, in our framework, language models only need to consider natural sequence changes [11], rather than the artificial mutations introduced in deep mutational scanning (DMS) experiments; evo-velocity therefore benefits by considering both the language model likelihood and semantic similarity [16]. Language models may provide successful evo-velocity predictions because their conditional likelihoods capture evolutionary contingency, which is a strong driver of natural sequence variation [47]. Our findings raise a number of interesting computational questions, including the degree to which the rules learned by language models are biologically interpretable (for example, in terms of thermostability or evolvability [28], [48]) and whether better protein language models could improve the performance and resolution of evo-velocity.

Promisingly, evo-velocity offers a new approach through which to reevaluate current evolutionary hypotheses. For example, when evaluating a potential hypothesis of eukaryote-to-prokaryote HGT among serpins [38], [39], evo-velocity instead predicted a more canonical evolutionary trajectory (**Figure 5**). While we mostly take a gene-centric approach to evolution [49], trajectories could also be integrated across multiple genes to provide insight into evolution at the level of pathways (as done for our analysis of glycolytic enzymes), gene modules, or even whole genomes. This might enable calibrating evo-velocity pseudotime to historical or geologic time, providing an additional method for dating evolutionary events. Evo-velocity also suggests a way to predict future evolution and to design novel protein sequences.

## Methods

### Language models

In this paper, we implement evo-velocity with masked language models, which are trained by masking certain residues in the input and predicting these residues in the output. For a sequence 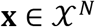, where 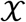 is the set of amino acids and *N* is the sequence length, the masked language modeling objective implicitly models a distribution over sequences through conditional likelihoods *p*(*x_i_*|**x**_[*N*]\{*i*}_) where **x**_[*N*]\{*i*}_ denotes the sequence without the residue at position *i*, sometimes referred to as the sequence context. Typically, these language models also learn a latent variable 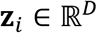 by learning a function 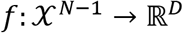 where 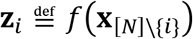 such that *p*(*x_i_*|**x**_[*n*]\{*i*}_, **z**_*i*_) = *p*(*x_i_*|**z**_*i*_).

We use two large-scale language models trained with a masked objective. We used the ESM-1b model [15] (obtained from https://github.com/facebookresearch/esm) trained on the March 2018 release of UniRef50 [17]. We also used the TAPE transformer model [14] (obtained from https://github.com/songlab-cal/tape) trained on the Pfam database release 32.0 [31]. Unless otherwise stated, we used ESM-1b as the default model for our experiments.

### Evo-velocity score computation

We compute an evo-velocity score that compares two sequences **x**^(*a*)^ and **x**^(*b*)^ as

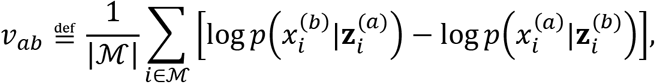

where 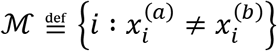 is the set of positions at which the amino acid residues disagree. We designed the evo-velocity score based on masked-language-model pseudolikelihoods [19] to efficiently approximate the change in likelihood of mutating sequence **x**^(*a*)^ to **x**^(*b*)^ and vice versa. The evo-velocity score is positive if moving from **x**^(*a*)^ to is more favorable and negative if moving from **x**^(*b*)^ to **x**^(*a*)^ is more favorable, so that *v_ab_* = –*v_ba_*.

In practice, **x**^(*a*)^ and **x**^(*b*)^ can disagree in length, so we first perform a global pairwise sequence alignment using the pairwise2 module in the Biopython Python package version 1.76 with a uniform substitution matrix and alignment parameters meant to discourage the introduction of sequence gaps (following the Biopython recommendations, we use a match score of 5, a mismatch penalty of −4, a gap-open penalty of −4, and a gap-extension penalty of −0.1). We ignore positions involving alignment gaps when computing the evo-velocity score, i.e., the evo-velocity score is only based on substitutions, since modeling the effect of an insertion or a deletion is less well defined when using a masked language model to predict mutations. We do not include gap characters when computing language model likelihoods.

### Constructing the sequence similarity network and evo-velocity transition matrix

To construct the sequence similarity network, we first use the language model to obtain a sequence embedding 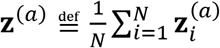 for each sequence **x**^(*a*)^ in the set of sequences-of-interest (for example, proteins within the same family) of size *M*. We use ESM-1b to compute the embeddings for each sequence as the 1,280-dimensional output of the last (i.e., the 33rd) hidden layer of the language model.

We then construct a directed graph where each node corresponds to a sequence and we connect a node to its *k*-nearest neighbors based on the Euclidean distance in the language model embedding space in 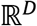. We can then use the evo-velocity scores and the KNN graph to construct a transition matrix 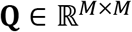, where

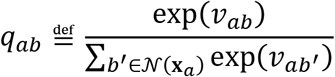

is the entry in the *a*th row and *b*th column of **Q** and 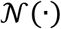 denotes the set of the neighbors in the KNN graph. Note that ∑_*b*∈[*M*]_ *q_ab_* = 1.

In all our experiments, we use the embedding function learned by the ESM-1b language model. To construct the KNN graph, we use the functionality provided by the Scanpy Python package version 1.6.1 [23]. In practice, higher values of *k* result in smoother, less noisy landscapes at the cost of higher computational effort. We find that values of *k* around 30 to 50 (our package defaults to 50) provide a good balance between robustness to noise and computational efficiency (though analyses involving less sequences overall or more homogeneous sequences can also tolerate lower values of *k* to speed up analysis); the 30-50 range has also shown good empirical performance in other KNN-based analyses that require robust estimation of the biological landscape [50]. In this paper, we use the values *k* = 30 for our cytochrome c and Spike experiments, *k* = 40 for our NP and Gag experiments, and *k* = 50 for our HA, globin, enolase, PGK, and serpin experiments.

### Network diffusion analysis and predicting roots

To find the root nodes, we can use the fixed points of a diffusion process based on the transition matrix **Q** [46], [51]. Given a diffusion probability vector **μ**^(*t*)^, we can find roots by running a diffusion process until a fixed point, i.e., **μ**^(∞)^ **= Q**^T^**μ**^(∞)^ (note that we take the transpose of the transition matrix to “reverse” the diffusion process, since our goal is to find the root nodes). We take the highest values of **μ**^(∞)^ to identify the root nodes, where we obtain **μ**^(∞)^ as the eigenvector of **Q**^T^ corresponding to an eigenvalue of 1. By default, we use a cutoff at the 98^th^ percentile of values in **μ**^(∞)^ to define the set of root nodes, as has been done previously [51]. We assume **Q** corresponds to a strongly connected directed graph, which is true if the KNN network consists of a single connected component (and which was true for all of our analyses); if the graph is strongly connected, then there is a unique value of **μ**^(∞)^ [46]. We scale the final values of the diffusion vector **μ**^(∞)^ to take values between 0 and 1, inclusive, and use the diffusion-based root estimation procedure implemented by the scVelo Python package version 0.2.2 [51].

### Diffusion pseudotime computation

We use diffusion pseudotime (DPT) to order sequences in evolutionary time. DPT is described in detail by Haghverdi et al. [26] and is closely related to the geodesic distance between two nodes in a graph. As done by Haghverdi et al., we denote the DPT score between a root node **x**^(root)^ and a node **x** as dpt(**x**^(root)^, **x**), which takes scaled values between 0 and 1, inclusive. We use the graph encoded by the transition matrix **Q**. Since the root-prediction analysis described above can yield potentially multiple roots, we define evo-velocity pseudotime as the average of DPT scores across the set of all root nodes 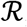, i.e.,

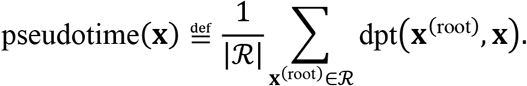

We use the DPT implementation provided by the Scanpy Python package.

### Plotting, data visualization, and statistical analysis

We used the UMAP algorithm [24] to visualize the KNN graph in two dimensions. All UMAP visualizations were obtained using the umap-learn Python package version 0.4.6 as wrapped by Scanpy. We generated boxplots using the seaborn Python package version 0.11.1; in all of our boxplots, the box extends from the first to third quartile, a horizontal line is drawn at the median, and whiskers extend to 1.5 times the interquartile range. We used the scipy version 1.4.1 Python package to compute correlations and statistical tests. A *P* value of less than 1 × 10^−308^ indicates a value that was below the floating-point precision of our computer.

### Embedding transfer

We can project evo-velocity, as encoded by the transition matrix **Q**, into an arbitrary embedding space (assuming that embeddings are available for all sequences) as done previously [51]. For a sequence **x**^(*a*)^ and **x**^(*b*)^, we denote the respective embeddings as **z**^(*a*)^ and **z**^(*b*)^. We then first compute the cosine-normalized translation vector separating sequences connected in the KNN graph, i.e.,

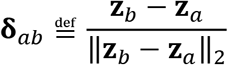

and we obtain the velocity projections as the expected displacement with respect to **Q**, i.e.,

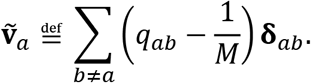

We use two main interpretable embedding spaces in our downstream analysis. The first is two-dimensional UMAP space, in which evo-velocity can be visualized as two-dimensional vectors. Once these vectors are computed, we use the streamplot and quiver plot functionality of the matplotlib Python package version 3.3.3 to visualize evo-velocity. The second interpretable embedding space we consider is one-hot-encoded sequence space, which we use to identify mutations that are associated with large changes in evo-velocity. To project evo-velocity into sequence space, we first construct a multiple sequence alignment of all *M* sequences using MAFFT version 7.475. A sequence **x** is then embedded into a one-hot-encoded vector 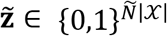, where 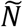 is the length of the alignment. The velocity projections take values in 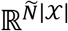, where we interpret each dimension as corresponding to a given residue in 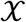 at a given site in 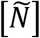.

### Deep mutational scan benchmarking

We obtained DMS values, all involving single-residue substitutions, and the corresponding DeepSequence [20] mutational effect predictions from Livesey and Marsh [18]. To compute mutational effect predictions for ESM-1b and TAPE, we used the evo-velocity score between the wildtype and mutant sequence as described above. As done by Livesey and March, we evaluated the performance of the mutational effect prediction as the absolute value of the Spearman correlation between the algorithm’s predicted mutational effect and the value reported by the original DMS study, restricting only to mutants considered by the original DMS studies. We used all DMS studies from Livesey and Marsh for which there were DeepSequence results available.

### UniRef50 sequence similarity computational control

We wanted to quantify if our evo-velocity results, including evo-velocity pseudotime, could be explained by sequence similarity to the training set. We obtained this training set from ftp://ftp.uniprot.org/pub/databases/uniprot/previous_releases/release-2018_03/uniref/. We identified representative sequences in UniRef50 by searching for the literal presence of the sequence within UniRef50 or by mapping the protein accession information to UniProt IDs, if available, and then mapping the UniProt IDs to the corresponding UniRef50 cluster representative. Then, for each sequence in our evo-velocity analysis, we computed the sequence similarity score to each representative sequence in UniRef50 and took the maximum of these scores. To compute the sequence similarity score, we used the similarity ratio implemented by the fuzzywuzzy Python package version 0.18.0, which is based on the Levenshtein distance between two sequences and is normalized to take values between 0% and 100%, inclusive.

To perform the control experiment, we filtered out sequences with 80% or less sequence similarity to the training set, thereby excluding sequences that are far from the sequences considered by ESM-1b. We then evaluated the Spearman correlation between the similarity scores and pseudotime, both in terms of the directionality of the correlation (e.g., a positive correlation indicates that similarity to UniRef50 could be explaining pseudotime) and also in terms of the change in this correlation compared to the correlation obtained on the full set of sequences (**Table S3**). We also evaluated the ability for the overall pseudotemporal patterns (for example, correlation with sampling time or ordering of taxonomic classes) to reproduce those found when analyzing the full set of sequences.

### TAPE reproducibility computational control

We also wanted to see how robust our evo-velocity results were to the language model used to estimate the mutational likelihoods. We therefore obtained the TAPE transformer model as described above. We performed the evo-velocity analysis by keeping the KNN graph structure the same as in the ESM-1b analysis but using the evo-velocity scores obtained by the TAPE likelihoods. All other downstream analyses, including root prediction and pseudotime computation, were also kept the same. We then evaluated the ability for the final pseudotime output to reproduce the output obtained by performing the same analysis except with ESM-1b velocities.

### Influenza A NP evo-velocity analysis

We obtained 3,304 unique NP sequences from the NIAID Influenza Research Database (https://www.fludb.org) [52]. We restricted our analysis to sequences that were sampled from human hosts. Metadata included the year the sequences were sampled and the influenza subtype of the original virus. We performed KNN graph construction, evo-velocity computation, root prediction, diffusion pseudotime estimation, and UMAP velocity projection as described previously.

We obtained an ordered phylogenetic path from Gong et al. [28] of H3N2-subtype NP evolution from 1968 to 2007. We computed the ESM-1b evo-velocity score comparing adjacent sequences along this path and plotted the cumulative sum of these scores versus the order in the path (**Figure 2D**). We also compared the improvement in evo-velocity of this path to that of simulated paths. To simulate paths across our evo-velocity landscape, we began at the same starting sequence, used the same number of steps as the path of Gong et al., and only considered paths that ended in the same cluster of sequences as the end sequence of Gong et al.’s path. We used the transition matrix **Q** to define the probability of moving from node to node and we performed 30,000 random walks.

We obtained a phylogenetic tree of all NP sequences considered in the evo-velocity analysis by first aligning sequences with MAFFT followed by approximate maximum-likelihood tree construction using FastTree version 2.1 using a JTT+CAT model. The midpoint-rooted tree was visualized using the iTOL web tool (https://itol.embl.de/) [53].

We also projected evo-velocity into one-hot-encoding space to compute a 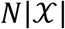-dimensional vector 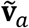 for each sequence as described previously; we then averaged these vectors across all sequences and inspected the top five mutations with the greatest magnitude change in the resulting average. We then located these mutations onto a reference sequence from 1934 H1N1 NP (UniProt ID: P03466), for which linear T-cell epitope data is available through the Immune Epitope Database (https://www.iedb.org/) [30]. We restricted our consideration to linear epitopes of influenza NP with positive validation in a T-cell assay.

We also conducted an ablation study to test the robustness of evo-velocity results when using simpler methods for computing sequence embeddings or evo-velocity scores. We recomputed the KNN graph based on 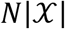-dimensional one-hot embeddings followed by dimensionality reduction based on the top-100 principal components to enable more efficient estimation of nearest-neighbor relationships. We recomputed evo-velocity scores based on the BLOSUM62 amino-acid substitution scores averaged across the set of differing positions, i.e., 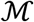, for each edge (obtained via global pairwise alignment with a uniform substitution matrix). We reran analysis using binary embeddings or BLOSUM62 velocities or both, while holding all other parts of the pipeline constant. As a negative control, we also computed velocities by sampling from a Gaussian distribution with zero mean and unit variance and reran analysis with all other parts of the pipeline constant.

### Influenza A HA evo-velocity analysis

We obtained 8,115 unique HA H1 sequences from the NIAID Influenza Research Database (https://www.fludb.org) [52]. We restricted our analysis to sequences that were sampled from human hosts. Metadata included the year the sequences were sampled and the influenza subtype of the original virus. We performed KNN graph construction, evo-velocity computation, root prediction, diffusion pseudotime estimation, and UMAP velocity projection as described previously.

### HIV-1 Gag evo-velocity analysis

We obtained 18,018 unique Gag sequences from the Los Alamos National Laboratory HIV sequence database (https://www.hiv.lanl.gov). Metadata included the year the sequences were sampled and the HIV subtype of the original virus. We performed KNN graph construction, evo-velocity computation, root prediction, diffusion pseudotime estimation, and UMAP velocity projection as described previously. We obtained four SIVcpz Gag sequences with high-quality, manual annotation from UniProt (https://www.uniprot.org/) [54]. These sequences were obtained from SIVcpz isolates MB66 (UniProt ID: Q1A268), EK505 (UniProt ID: Q1A250), TAN1 (UniProt ID: Q8AII2), and GAB1 (UniProt ID: P17282).

### SARS-CoV-2 Spike evo-velocity analysis

We obtained 46,986 unique, full-length Spike sequences from the May 27, 2021 GISAID release (https://www.gisaid.org/) [55]. Metadata included the date the sequences were sampled. We performed KNN graph construction, evo-velocity computation, root prediction, diffusion pseudotime estimation, and UMAP velocity projection as described previously. We determined the location of clusters corresponding to known variants-of-concern based on known marker mutations including D614G, N501Y (for B.1.1.7, B.1.351, and P.1), K417N (for B.1.351), P681H (for B.1.1.7), E154K (for B.1.617.1), and T478K (for B.1.617.2) [34].

### Globins evo-velocity analysis

We obtained 6,097 globin sequences from UniProt. We restricted our analysis to eukaryotic sequences within the “globin” family and to sequences between 135 and 155 residues in length, inclusive, which was done based on a clear mode in the distribution of sequence lengths and was meant to preserve mostly homologous sequences in our analysis. Metadata included the taxonomic lineage of each sequence and, for some of the sequences, annotations indicating the type of globin. We performed KNN graph construction, evo-velocity computation, root prediction, diffusion pseudotime estimation, and UMAP velocity projection as described previously. We obtained the rooted phylogenetic tree of globins and the inferred ancestral sequences from Pillai et al. [35].

### Cytochrome c evo-velocity analysis

We obtained 2,128 cytochrome c sequences from UniProt. We restricted our analysis to eukaryotic sequences within the “cytochrome c” family and to sequences between 100 and 115 residues in length, inclusive, which was done based on a clear mode in the distribution of sequence lengths and was meant to preserve mostly homologous sequences in our analysis. Metadata included the taxonomic lineage of each sequence. We performed KNN graph construction, evo-velocity computation, root prediction, diffusion pseudotime estimation, and UMAP velocity projection as described previously. We obtained the approximate dates and geologic eons of the emergences of different organisms from Hedges et al. [37].

### Enolase evo-velocity analysis

We obtained 31,901 enolase sequences from UniProt. We restricted our analysis to sequences within the “enolase” family and to sequences between 412 and 448 residues in length, inclusive, which was done based on a clear mode in the distribution of sequence lengths and was meant to preserve mostly homologous sequences in our analysis. Metadata included the taxonomic lineage of each sequence. We performed KNN graph construction, evo-velocity computation, root prediction, diffusion pseudotime estimation, and UMAP velocity projection as described previously.

We obtained unrooted phylogenetic trees of enolase based on the subset of our UniProt sequences with high-quality, manual annotation. We then performed a multiple sequence alignment with MAFFT and performed phylogenetic reconstruction on the alignment with PhyML version 3.3.20200621 using a JTT model with gamma-distributed among-site rate variation and empirical state frequencies [56]. The unrooted tree was visualized using the iTOL web tool.

### PGK evo-velocity analysis

We obtained 30,455 PGK sequences from UniProt. We restricted our analysis to sequences within the “phosphoglycerate kinase” family and to sequences between 385 and 420 residues in length, inclusive, which was done based on a clear mode in the distribution of sequence lengths and was meant to preserve mostly homologous sequences in our analysis. Metadata included the taxonomic lineage of each sequence. We performed KNN graph construction, evo-velocity computation, root prediction, diffusion pseudotime estimation, and UMAP velocity projection as described previously.

We obtained unrooted phylogenetic trees of enolase based on the subset of our UniProt sequences with high-quality, manual annotation. We then performed a multiple sequence alignment with MAFFT and performed phylogenetic reconstruction on the alignment with PhyML using a JTT model with gamma-distributed among-site rate variation and empirical state frequencies. The unrooted tree was visualized using the iTOL web tool.

### Serpins evo-velocity analysis

We obtained 22,737 serpin sequences from UniProt. We restricted our analysis to sequences within the “serpin” family and to sequences between 300 and 525 residues in length, inclusive, which was done based on a clear mode in the distribution of sequence lengths and was meant to preserve mostly homologous sequences in our analysis. Metadata included the taxonomic lineage of each sequence. We performed KNN graph construction, evo-velocity computation, root prediction, diffusion pseudotime estimation, and UMAP velocity projection as described previously.

We obtained unrooted phylogenetic trees of enolase based on the subset of our UniProt sequences with high-quality, manual annotation. We then performed a multiple sequence alignment with MAFFT and performed phylogenetic reconstruction on the alignment with PhyML using a JTT model with gamma-distributed among-site rate variation and empirical state frequencies. The unrooted tree was visualized using the iTOL web tool.

## Supporting information

Data S1: DMS benchmark results

## Data and code availability

Data used in our analysis has been deposited to Zenodo at doi:10.5281/zenodo.4891758. Code used in our analysis has been deposited to Zenodo at doi:10.5281/zenodo.4891819. Our code and links to data are also available on GitHub at https://github.com/brianhie/evolocity.

## Acknowledgements

We thank Nicholas Bhattacharya, Anne Dekas, Bennett Kapili, Michael Kim, Adam Lerer, Hanon McShea, Joshua Meier, Paula Welander, Ellen Zhong, and members of the Peter Kim Laboratory for helpful comments and discussion. We thank the Stanford Research Computing Center for providing computational resources and support through the Sherlock cluster. B.L.H. acknowledges the support of the Stanford Science Fellows program. This work was supported by the Chan Zuckerberg Biohub (P.S.K.).

## Author contributions

All authors were involved in project conceptualization and investigation. B.L.H. wrote the software, performed the computational experiments, and wrote the initial paper draft. All authors interpreted the results and wrote the final paper.

## Competing interests

The authors declare no competing interests.

## Supplemental data

**Data S1:** Mutational effect prediction benchmarking results (CSV file is included).

## Supplementary figures and figure captions

**Figure S1:**
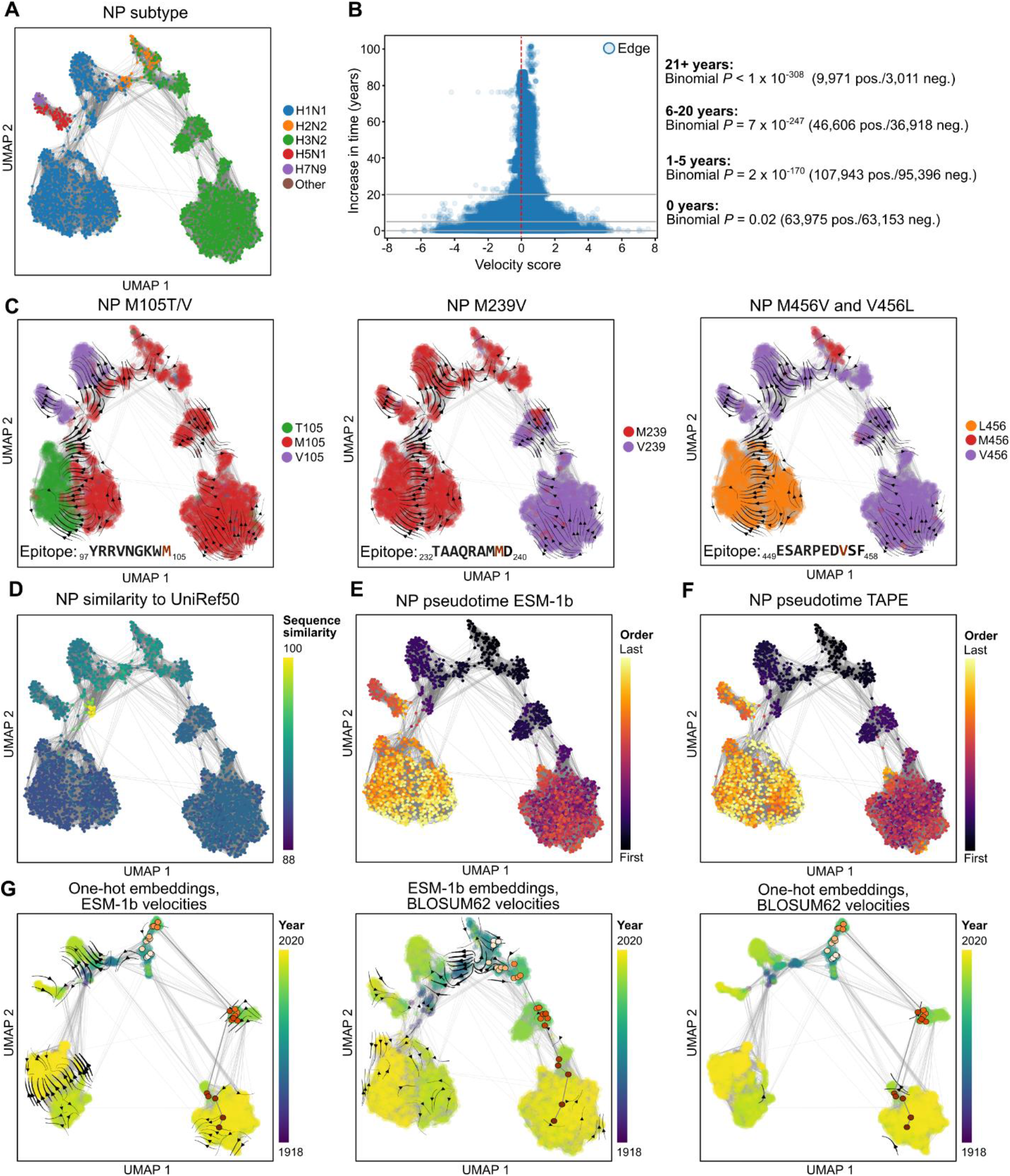
Additional figures for nucleoprotein evo-velocity analysis. (**A**) The NP sequence landscape shows structure corresponding to influenza subtype. (**B**) By stratifying edges based on the sampling time difference between their two corresponding sequences and quantifying bias toward positive or negative evo-velocity scores using a binomial test, we found that the bias toward positive evo-velocity scores increases as time increases. (**C**) Mutations with strong magnitude changes in evo-velocity are also located in experimentally-validated T-cell epitopes (**Table S2**). (**D**) All NP sequences belong to a single UniRef50 cluster, which has as its representative a sequence from 1934 H1N1. (**E, F**) Evo-velocity pseudotime of NP based on ESM-1b- or TAPE-based evo-velocity scores have high correlation (**Table S4**). (**G**) Replacing ESM-1b embeddings with one-hot sequence embeddings removes some of the known evolutionary continuity relationships from the visualization, especially in the well-studied trajectory of H3N2 NP evolution. Replacing ESM-1b evo-velocity scores with BLOSUM62 scores results in much weaker and more ambiguous evo-velocity flows when visualized.

**Figure S2:**
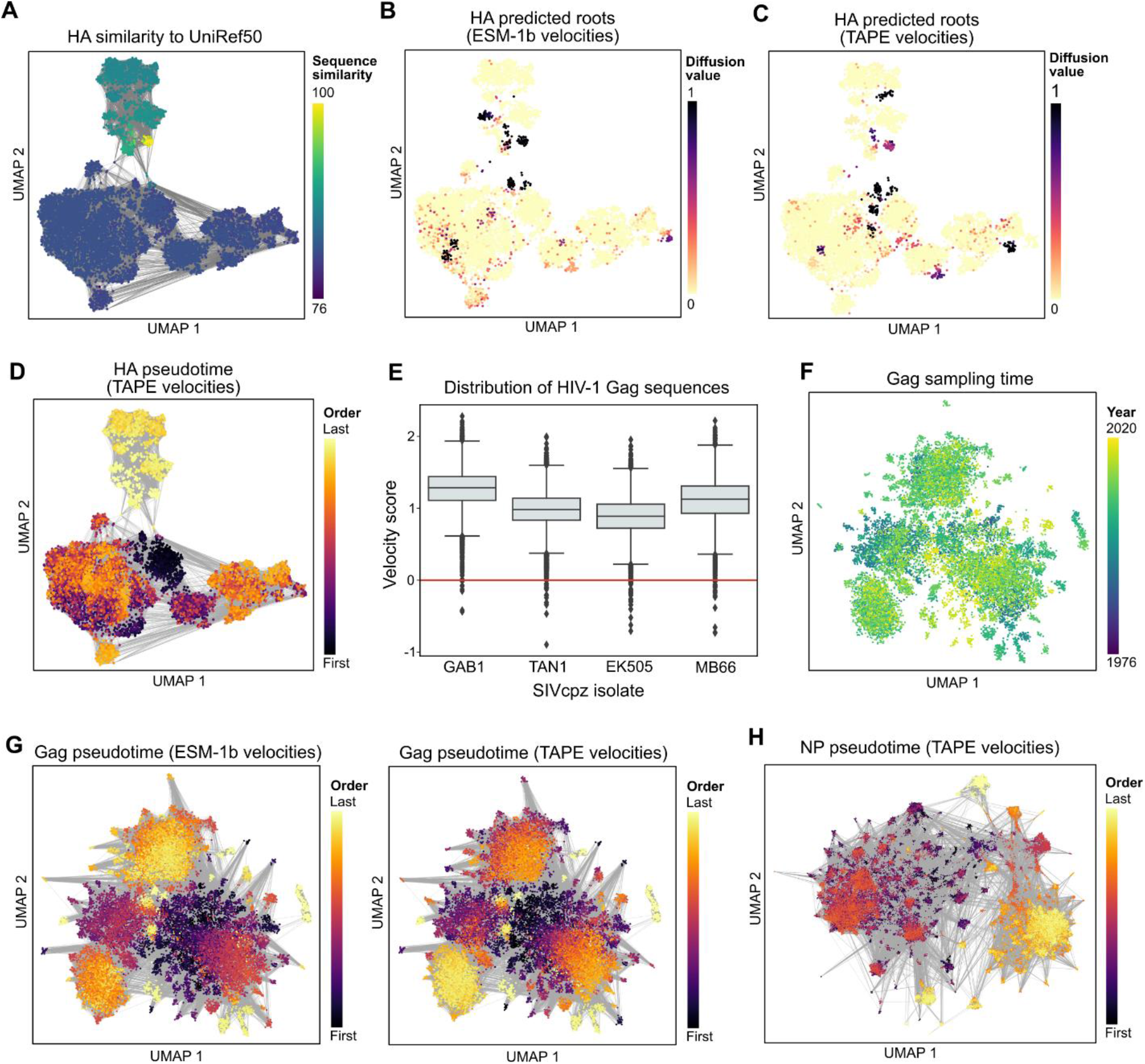
Additional figures for viral protein evo-velocity analyses. (**A**) Influenza A HA H1 sequences map to a single UniRef50 cluster, where the representative sequence is from a 1934 H1N1 strain. (**B**) Using ESM-1b-based evo-velocity scores, the inferred roots correspond to early twentieth-century H1N1 sequences, including 1918 influenza, as well as twenty-first-century 2009 H1N1 pandemic influenza. (**C, D**) In contrast, with TAPE-based velocities, the 2009 pandemic roots are identified but not the earlier 1918 pandemic roots, leading to evo-velocity pseudotimes that are higher for twentieth-century influenza. (**E**) Each boxplot visualizes the distribution of velocity scores for all HIV-1 Gag sequences in our analysis compared to a SIVcpz Gag sequence from a given isolate. On average, HIV-1 Gag sequences have strong positive evo-velocity scores compared to the four SIVcpz Gag sequences. Box extends from first to third quartile with line at the median, whiskers extend to 1.5 times the interquartile range, and diamonds indicate outlier points. (**F**) There is less temporal structure in the sequence landscape of HIV-1 Gag, reflecting the lack of immune pressure on HIV-1. (**G**) Evo-velocity pseudotime of Gag based on ESM-1b- or TAPE-based evo-velocity scores have high correlation (**Table S4**). (**H**) Evo-velocity pseudotime of SARS-CoV-2 Spike based on ESM-1b- or TAPE-based evo-velocity scores have high correlation (**Table S4**); compare to **Figure 3G**.

**Figure S3:**
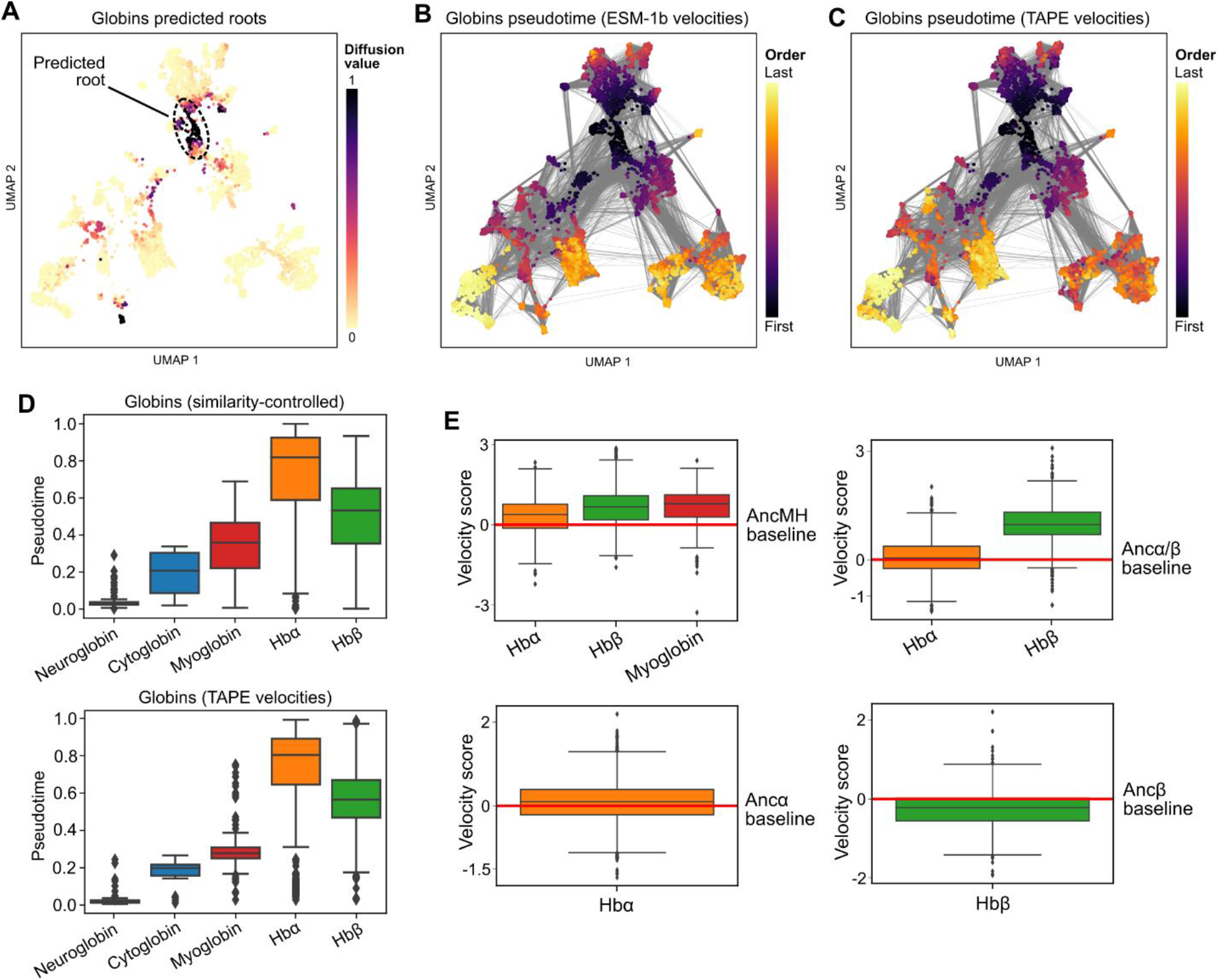
Additional figures for globin evo-velocity analysis. (**A**) The main root region predicted for globin evolution is closest to (and includes) neuroglobin (**Figure 4B**). (**B**) Evo-velocity pseudotime is therefore lowest for neuroglobin and increases radiating outward from that portion of the graph, with Hbα and Hbβ predicted to be most recent in pseudotime. (**C**) TAPE-based evo-velocity scores lead to pseudotime values that strongly correlate with those based on ESM-1b evo-velocity scores (**Table S4**). (**D**) Pseudotemporal relationships when controlling for similarity to UniRef50 or when using TAPE-based evo-velocity computation reproduce those in our main analysis; compare to **Figure 4C**. (**E**) Extant Hbα, Hbβ, and myoglobin sequences have positive evo-velocity scores, on average, compared to a reconstructed myoglobin/hemoglobin ancestor (AncMH) as the baseline sequence, consistent with AncMH preceding extant globins in evolutionary time. Extant Hbβ sequences also have positive velocities with respect to a reconstructed Hbα/Hbβ ancestor (Ancα/β), but this is not observed for extant Hbα sequences, predicting that extant Hbαs are more similar to Ancα/β than extant Hbβs and corroborated by the phylogeny of Pillai et al. [35] (**Figure 4A**). Evo-velocity also predicts extant Hbαs and Hbβs show little improvement in evo-velocity from their respective most proximal ancestors. Together, these results are consistent with evo-velocity scores increasing over greater stretches of evolutionary time. For all boxplots: box extends from first to third quartile with line at the median, whiskers extend to 1.5 times the interquartile range, and diamonds indicate outlier points.

**Figure S4:**
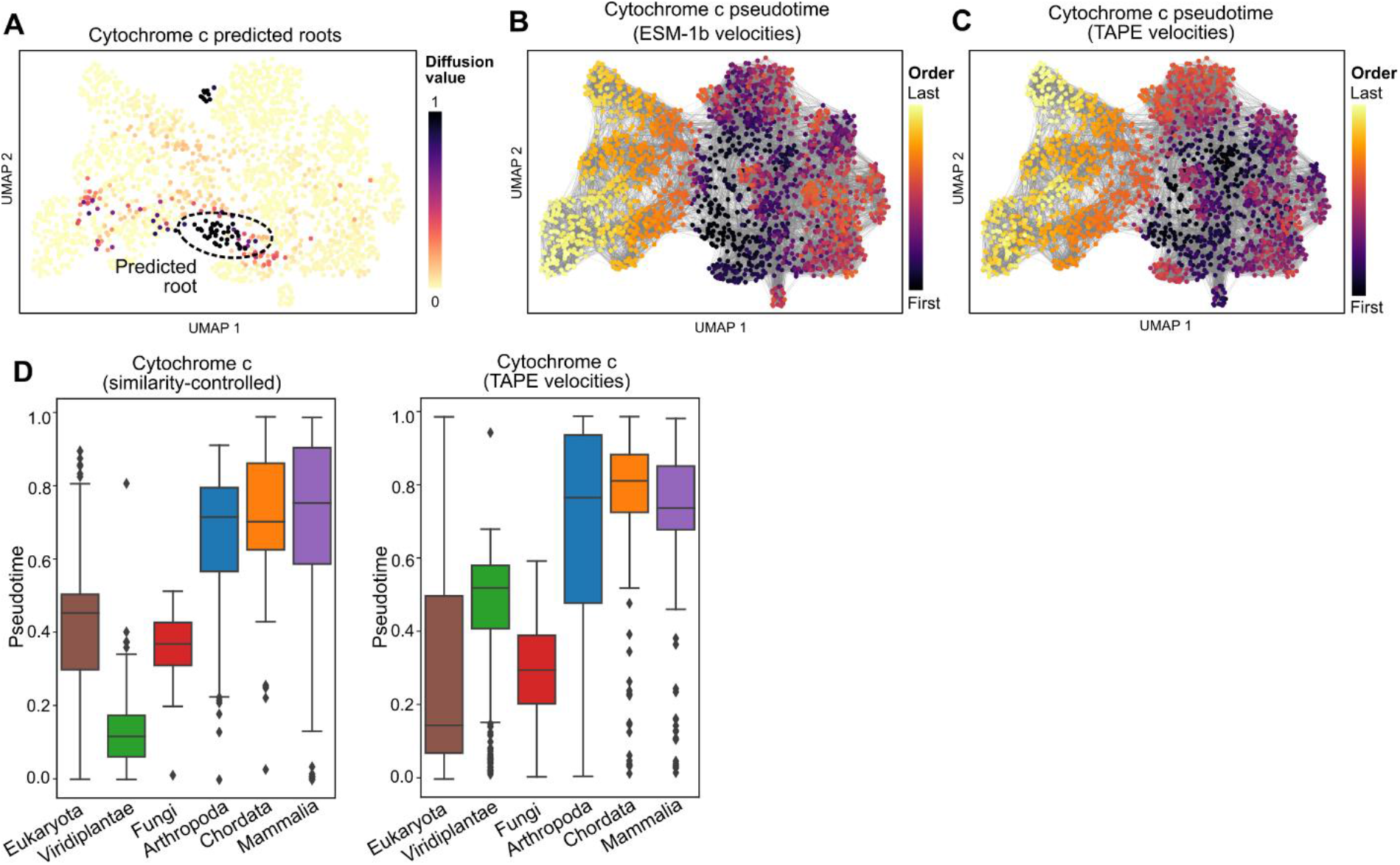
Additional figures for evo-velocity analysis of cytochrome c. (**A**) Evo-velocity predicts root regions in the extant sequence landscape among single-celled eukaryotes and green algae. (**B, C**) Evo-velocity pseudotime of cytochrome c based on ESM-1b or TAPE velocities have high correlation (**Table S4**). (**D**) Pseudotemporal relationships when controlling for similarity to UniRef50 or when using TAPE-based evo-velocity computation largely reproduce those in our main analysis (compare to **Figure 4E**) especially when comparing the “lower-order” and “higher-order” taxonomic labels, although TAPE places viridiplantae after fungi in pseudotime and filtering based on sequence similarity to UniRef50 removes many of the earliest eukaryotes in pseudotime when analyzing the full dataset. Box extends from first to third quartile with line at the median, whiskers extend to 1.5 times the interquartile range, and diamonds indicate outlier points.

**Figure S5:**
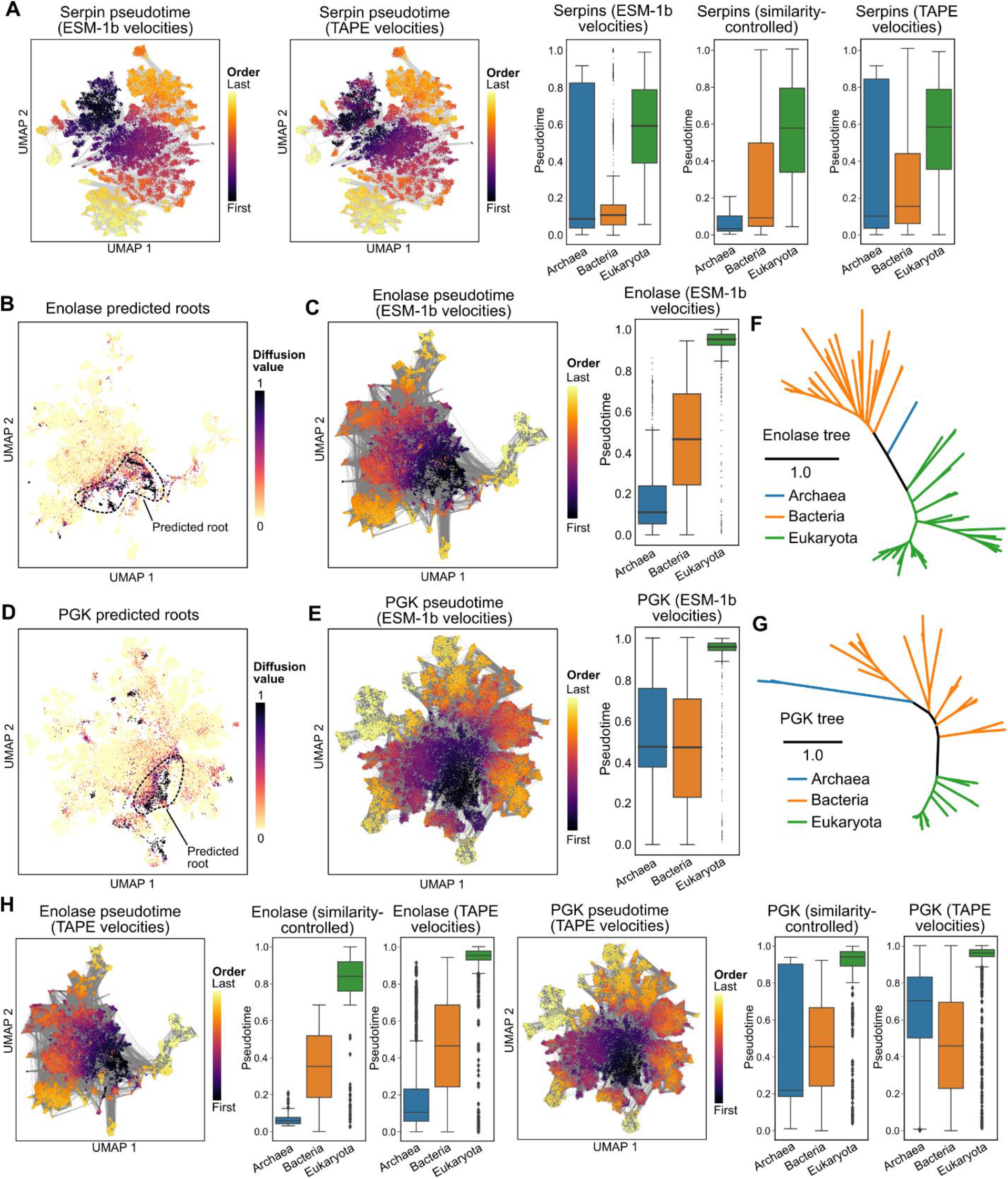
Additional figures for highly conserved protein evo-velocity analyses. (**A**) For the family of serpins, pseudotemporal orderings of the three domains of life were reproducible when using TAPE-based evo-velocity computation and when filtering based on similarity to UniRef50. In all cases, prokaryotes precede eukaryotes in evo-velocity pseudotime. (**B, C**) Enolase is predicted to be rooted in a region with archaeal and some bacterial sequences, with eukaryota occurring last in evo-velocity pseudotime. (**D, E**) PGK is predicted to be rooted in a region with archaeal and bacterial sequences, with eukaryota occurring last in evo-velocity pseudotime. (**F**) The unrooted phylogenetic tree of manually curated enolase sequences shows archaeal sequences as more proximal to the eukaryota than bacterial sequences. (**G**) In contrast, the unrooted phylogenetic tree of manually curated PGK sequences shows bacterial sequences as more proximal to the eukaryota than archaeal sequences. (**H**) For both enolase and PGK, pseudotemporal orderings of the three domains of life were reproducible when using TAPE-based evo-velocity computation and when filtering based on similarity to UniRef50 (compare to **C** and **E**). For all boxplots: box extends from first to third quartile with line at the median, whiskers extend to 1.5 times the interquartile range, and diamonds indicate outlier points.

## Supplementary Tables

**Table S1:** Taxonomic composition of UniRef50. The number of sequences in UniRef50 that belong to different taxonomic categories. Most sequences in UniRef50 are bacterial, though we note that ESM-1b had no access to these taxonomic labels at training time.

| Taxonomy |  | # Sequences | % |
| --- | --- | --- | --- |
| Archaea |  | 776,374 | 2.57% |
| Bacteria |  | 18,032,582 | 59.79% |
| Eukaryota | Primate | 160,932 | 0.53% |
|  | Other mammalia | 341,837 | 1.13% |
|  | Other chordata | 950,939 | 3.15% |
|  | Arthropoda | 1,521,727 | 5.05% |
|  | Viridiplantae | 2,037,089 | 6.75% |
|  | Fungi | 2,880,452 | 9.55% |
|  | Other eukaryotes | 2,783,754 | 9.23% |
| Metagenome |  | 637,280 | 2.11% |
| Other/unclassified |  | 39,121 | 0.13% |
| Total |  | 30,162,087 | 100% |

**Table S2:** Top five mutations by evo-velocity rank and corresponding IEDB epitopes. Mutations were ranked by the magnitude of the average evo-velocity vector obtained by projecting the velocities into sequence space (**Methods**) and the top five were further investigated for location in T-cell epitopes. All involve single-nucleotide mutations from a methionine to a hydrophobic or a polar-uncharged amino acid residue. Also see **Figures 2G** and **S1C**.

| Rank | Mutations | Epitope | # publications in IEDB |
| --- | --- | --- | --- |
| 1 | M105V | YRRVNGKWM | 5 |
| 2 | M374I | ASNENMETM | 105 |
| 3 | M481I, M481V | SPIVPSFDM | 5 |
| 4 | M239V | TAAQRAMMD | 3 |
| 5 | M456V, V456L | ESARPEDVSF | 6 |

**Table S3:**
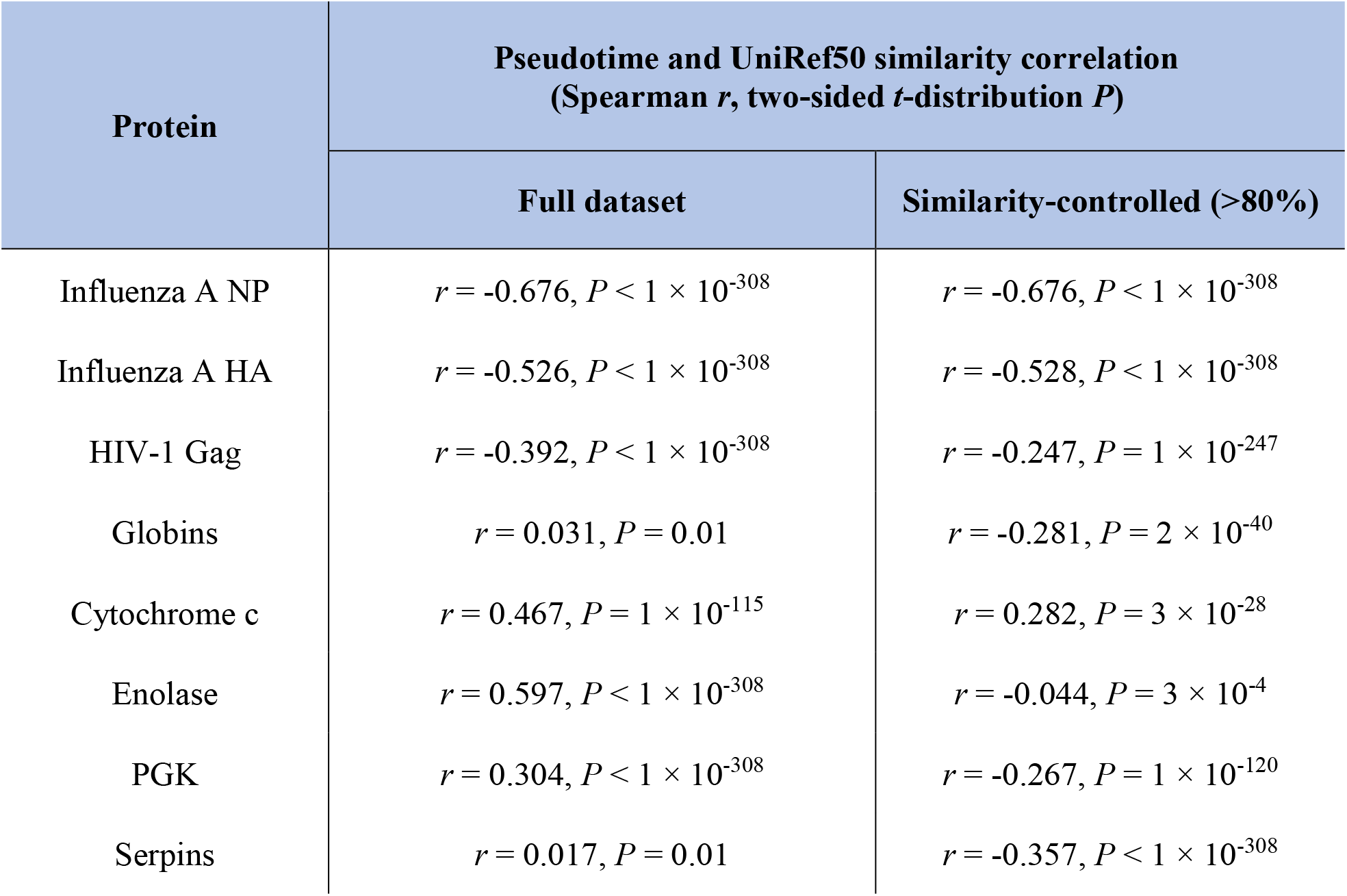
Correlation between evo-velocity pseudotime and sequence similarity to UniRef50. There is no consistent pattern in the directionality of the correlation between evo-velocity pseudotime and sequence similarity to UniRef50, indicating that sequence similarity does not trivially explain pseudotime. “Full dataset” indicates the results from analyzing all sequences while “similarity-controlled” indicates the results from restricting analysis to the sequences with greater than 80% sequence similarity to UniRef50 (**Methods**). In this latter setting, for all proteins, we were able to reproduce the results obtained from running evo-velocity on the full dataset.

| Protein | Pseudotime and UniRef50 similarity correlation<br>(Spearman $r$, two-sided $t$-distribution $P$) | |
| --- | --- | --- |
|  | Full dataset | Similarity-controlled (>80%) |
| Influenza A NP | $r = -0.676, P < 1 \times 10^{-308}$ | $r = -0.676, P < 1 \times 10^{-308}$ |
| Influenza A HA | $r = -0.526, P < 1 \times 10^{-308}$ | $r = -0.528, P < 1 \times 10^{-308}$ |
| HIV-1 Gag | $r = -0.392, P < 1 \times 10^{-308}$ | $r = -0.247, P = 1 \times 10^{-247}$ |
| Globins | $r = 0.031, P = 0.01$ | $r = -0.281, P = 2 \times 10^{-40}$ |
| Cytochrome c | $r = 0.467, P = 1 \times 10^{-115}$ | $r = 0.282, P = 3 \times 10^{-28}$ |
| Enolase | $r = 0.597, P < 1 \times 10^{-308}$ | $r = -0.044, P = 3 \times 10^{-4}$ |
| PGK | $r = 0.304, P < 1 \times 10^{-308}$ | $r = -0.267, P = 1 \times 10^{-120}$ |
| Serpins | $r = 0.017, P = 0.01$ | $r = -0.357, P < 1 \times 10^{-308}$ |

**Table S4:** Pseudotime reproducibility with TAPE velocities. The correlation between computed pseudotime using ESM-1b or TAPE to determine the evo-velocity scores. Cells shaded in light blue indicate correlations greater than 0.8. HA pseudotimes were not correlated between ESM-1b and TAPE due to the inability of TAPE to identify roots among the twentieth-century trajectory of HA evolution (**Figure S2B-D**). All other proteins had strong pseudotime reproducibility between the two language models.

| Protein | ESM-1b and TAPE pseudotime correlation<br>(Spearman $r$, two-sided $t$-distribution $P$) |
| --- | --- |
| Influenza A NP | $r = 0.926, P < 1 \times 10^{-308}$ |
| Influenza A HA | $r = -0.028, P = 0.01$ |
| HIV-1 Gag | $r = 0.814, P < 1 \times 10^{-308}$ |
| SARS-CoV-2 Spike | $r = 0.902, P < 1 \times 10^{-308}$ |
| Globins | $r = 0.893, P < 1 \times 10^{-308}$ |
| Cytochrome c | $r = 0.811, P < 1 \times 10^{-308}$ |
| Enolase | $r = 0.932, P < 1 \times 10^{-308}$ |
| PGK | $r = 0.948, P < 1 \times 10^{-308}$ |
| Serpins | $r = 0.955, P < 1 \times 10^{-308}$ |

**Table S5:** Evo-velocity ablation results for influenza A NP. We obtained comparable, if slightly weaker, pseudotime correlation when either using binary sequence embeddings to construct the KNN graph or using BLOSUM62 scores to compute velocities (or both). Replacing velocity scores with random, Gaussian noise resulted in loss of correlation between pseudotime and sampling year. PCA: Principal component analysis.

| Embedding | Velocity | Correlation with ESM-1b embeddings and velocities | Correlation with sampling year |
| --- | --- | --- | --- |
| ESM-1b | ESM-1b | N/A | 0.49 |
| One-hot + PCA | ESM-1b | 0.89 | 0.47 |
| ESM-1b | BLOSUM62 | 0.80 | 0.40 |
| One-hot + PCA | BLOSUM62 | 0.51 | 0.31 |
| ESM-1b | Random | -0.09 | -0.01 |

**Table S6:** Correlation between pseudotime and sequence length. We observed no consistent pattern in the correlation between pseudotime and the length of sequences, suggesting that differing sequences lengths across a landscape does not explain evo-velocity patterns.

| Protein | Pseudotime and sequence length correlation<br>(Spearman $r$, two-sided $t$-distribution $P$) |
| --- | --- |
| Influenza A NP | $r = 0.030, P = 0.08$ |
| Influenza A HA | $r = 0.534, P < 1 \times 10^{-308}$ |
| HIV-1 Gag | $r = -0.352, P < 1 \times 10^{-308}$ |
| SARS-CoV-2 Spike | $r = -0.681, P < 1 \times 10^{-308}$ |
| Globins | $r = -0.166, P = 4 \times 10^{-39}$ |
| Cytochrome c | $r = -0.414, P = 6 \times 10^{-89}$ |
| Enolase | $r = 0.273, P < 1 \times 10^{-308}$ |
| PGK | $r = 0.099, P = 2 \times 10^{-67}$ |
| Serpins | $r = -0.087, P = 4 \times 10^{-39}$ |

## Notes

https://github.com/brianhie/evolocity

### Competing Interest Statement

The authors have declared no competing interest.

## References

[1] C. Darwin, On the Origin of Species. 1909.

[2] J. Maynard Smith, “Natural selection and the concept of a protein space,”Nature, vol. 225, no. 5232, pp. 563–564, 1970.

[3] J. A. G. M. De Visser and J. Krug, “Empirical fitness landscapes and the predictability of evolution,”Nature Reviews Genetics, vol. 15, no. 7. pp. 480–490, 2014.

[4] M. Lässig, V. Mustonen, and A. M. Walczak, “Predicting evolution,”Nat. Ecol. Evol.,vol. 1, no. 3, pp. 1–9, 2017.

[5] M. Łuksza and M. Lässig, “A predictive fitness model for influenza,”Nature, vol. 507, no. 7490, pp. 57–61, 2014.

[6] M. C. Weiss et al., “The physiology and habitat of the last universal common ancestor,”Nat. Microbiol., vol. 1, no. 9, pp. 1–8, 2016.

[7] S. Guindon and O. Gascuel, “A Simple, Fast, and Accurate Algorithm to Estimate Large Phylogenies by Maximum Likelihood,”Syst. Biol., vol. 52, no. 5, pp. 696–704, 2003.

[8] J. P. Huelsenbeck, J. P. Bollback, and A. M. Levine, “Inferring the root of a phylogenetic tree,”Syst. Biol., vol. 51, no. 1, pp. 32–43, 2002.

[9] Y. Kim, F. Koehler, A. Moitra, E. Mossel, and G. Ramnarayan, “How Many Subpopulations Is Too Many? Exponential Lower Bounds for Inferring Population Histories,”J. Comput. Biol., vol. 27, no. 4, pp. 136–157, 2020.

[10] S. Wright, “The roles of mutation, inbreeding, crossbreeding and selection in evolution,” Sixth Int. Congr. Genet., vol. 1, no. 6, pp. 355–366, 1932.

[11] D. M. Weinreich, N. F. Delaney, M. A. DePristo, and D. L. Hartl, “Darwinian evolution can follow only very few mutational paths to fitter proteins,”Science, vol. 312, no. 5770, pp. 111–114, 2006.

[12] R. Dawkins, Climbing Mount Improbable. 1997.

[13] T. Bepler and B. Berger, “Learning protein sequence embeddings using information from structure,” in 7th International Conference on Learning Representations, 2019, arXiv, cs.LG:1902.08661.

[14] R. Rao et al., “Evaluating Protein Transfer Learning with TAPE,”Adv. Neural Inf. Process. Syst., vol. 32, pp. 9686–9698, 2019.

[15] A. Rives et al., “Biological structure and function emerge from scaling unsupervised learning to 250 million protein sequences,”Proc. Natl. Acad. Sci., vol. 118, no. 15, p. e2016239118, 2021.

[16] B. Hie, E. Zhong, B. Berger, and B. Bryson, “Learning the language of viral evolution and escape,”Science, vol. 371, no. 6526, pp. 284–288, 2021.

[17] B. E. Suzek, H. Huang, P. McGarvey, R. Mazumder, and C. H. Wu, “UniRef: Comprehensive and non-redundant UniProt reference clusters,”Bioinformatics, vol. 23, no. 10, pp. 1282–1288, 2007.

[18] B. J. Livesey and J. A. Marsh, “Using deep mutational scanning to benchmark variant effect predictors and identify disease mutations,”Mol. Syst. Biol., vol. 16, no. 7, p. e9380, 2020.

[19] C. Hsu, H. Nisonoff, C. Fannjiang, and J. Listgarten, “Combining evolutionary and assay-labelled data for protein fitness prediction,”bioRxiv, 10.1101/2021.03.28.437402, 2021.

[20] A. J. Riesselman, J. B. Ingraham, and D. S. Marks, “Deep generative models of genetic variation capture the effects of mutations,”Nat. Methods, vol. 15, no. 10, pp. 816–822, 2018.

[21] Y. W. Yu, N. M. Daniels, D. C. Danko, and B. Berger, “Entropy-Scaling Search of Massive Biological Data,” Cell Syst., vol. 1, no. 2, pp. 130–140, 2015.

[22] D. M. Mccandlish, “Visualizing fitness landscapes,”Evolution, vol. 65, no. 6, pp. 1544–1558, 2011.

[23] F. A. Wolf, P. Angerer, and F. J. Theis, “SCANPY: Large-scale single-cell gene expression data analysis,”Genome Biol., vol. 19, no. 1, p. 15, 2018.

[24] L. McInnes and J. Healy, “UMAP: Uniform Manifold Approximation and Projection for Dimension Reduction,” arXiv, stat.ML:1802.03426, 2018.

[25] A. L. Barabási, N. Gulbahce, and J. Loscalzo, “Network medicine: A network-based approach to human disease,”Nature Reviews Genetics, vol. 12, no. 1. pp. 56–68, 2011.

[26] L. Haghverdi, M. Büttner, F. A. Wolf, F. Buettner, and F. J. Theis, “Diffusion pseudotime robustly reconstructs lineage branching,”Nat. Methods, vol. 13, pp. 845–848, 2016.

[27] G. La Manno et al.,“RNA velocity of single cells,” Nature, vol. 560, no. 7719, pp. 494–498, 2018.

[28] L. I. Gong, M. A. Suchard, and J. D. Bloom, “Stability-mediated epistasis constrains the evolution of an influenza protein,”eLife, vol. 2013, no. 2, p. e00631, 2013.

[29] T. C. Sutton, “The pandemic threat of emerging H5 and H7 avian influenza viruses,”Viruses, vol. 10, no. 9. p. 461, 2018.

[30] R. Vita et al.,“The immune epitope database (IEDB) 3.0,” Nucleic Acids Res., vol. 43, no. D1, pp. D405–D412, 2015.

[31] S. El-Gebali et al.,“The Pfam protein families database in 2019,” Nucleic Acids Res., vol. 47, no. D1, 2019.

[32] D. M. Eckert and P. S. Kim, “Mechanisms of Viral Membrane Fusion and Its Inhibition,” Annu. Rev. Biochem., vol. 70, no. 1, pp. 777–810, 2001.

[33] P. M. Sharp and B. H. Hahn, “Origins of HIV and the AIDS pandemic,”Cold Spring Harb. Perspect. Med., vol. 1, no. 1, p. a006841, 2011.

[34] R. P. Walensky, H. T. Walke, and A. S. Fauci, “SARS-CoV-2 Variants of Concern in the United States-Challenges and Opportunities,”JAMA - Journal of the American Medical Association, vol. 325, no. 11. pp. 1037–1038, 2021.

[35] A. S. Pillai et al.,“Origin of complexity in haemoglobin evolution,” Nature, vol. 581, no. 7809, pp. 480–485, 2020.

[36] P. J. McLaughlin and M. O. Dayhoff, “Eukaryote evolution: A view based on cytochrome c sequence data,”J. Mol. Evol., vol. 2, no. 2-3, pp. 99–116, 1973.

[37] S. B. Hedges, J. Marin, M. Suleski, M. Paymer, and S. Kumar, “Tree of life reveals clock-like speciation and diversification,”Mol. Biol. Evol., vol. 32, no. 4, pp. 835–845, 2015.

[38] J. A. Irving, P. J. M. Steenbakkers, A. M. Lesk, H. J. M. Op den Camp, R. N. Pike, and J. C. Whisstock, “Serpins in prokaryotes,”Mol. Biol. Evol., vol. 19, no. 11, pp. 1881–1890, 2002.

[39] T. H. Roberts, J. Hejgaard, N. F. W. Saunders, R. Cavicchioli, and P. M. G. Curmi, “Serpins in unicellular Eukarya, Archaea, and Bacteria: Sequence analysis and evolution,”J. Mol. Evol., vol. 59, no. 4, pp. 437–447, 2004.

[40] M. A. Spence, M. D. Mortimer, A. M. Buckle, B. Q. Minh, and C. J. Jackson, “A comprehensive phylogenetic analysis of the serpin superfamily,”Mol. Biol. Evol., p. msab081, 2021.

[41] S. Potter and L. A. Fothergill-Gilmore, “Molecular evolution: The origin of glycolysis,”Biochem. Educ., vol. 21, no. 1, pp. 45–48, 1993.

[42] M. Piast, I. Kustrzeba-Wójcicka, M. Matusiewicz, and T. Banaś, “Molecular evolution of enolase,”Acta Biochim. Pol., vol. 52, no. 2, pp. 507–513, 2005.

[43] M. Rojas-Pirela et al.,“Phosphoglycerate kinase: Structural aspects and functions, with special emphasis on the enzyme from Kinetoplastea: Phosphoglycerate Kinase,”Open Biology, vol. 10, no. 11. p. 200302, 2020.

[44] S. J. Gould, Wonderful Life: The Burgess Shale and the Nature of History. WW Norton & Company, 1990.

[45] S. C. Morris, Life’s solution: Inevitable humans in a lonely universe. 2003.

[46] N. Masuda, M. A. Porter, and R. Lambiotte, “Random walks and diffusion on networks,”Physics Reports, vol. 716-717. pp. 1–58, 2017.

[47] V. C. Xie, J. Pu, B. P. Metzger, J. W. Thornton, and B. C. Dickinson, “Contingency and chance erase necessity in the experimental evolution of ancestral proteins,”eLife, vol. 10, p. e67336, 2021.

[48] J. D. Bloom, S. T. Labthavikul, C. R. Otey, and F. H. Arnold, “Protein stability promotes evolvability,”Proc. Natl. Acad. Sci. U. S. A., vol. 103, no. 15, pp. 5869–5874, 2006.

[49] R. Dawkins, The Selfish Gene. 1976.

[50] A. Narayan, B. Berger, and H. Cho, “Assessing single-cell transcriptomic variability through density-preserving data visualization,”Nat. Biotechnol., 2021.

[51] V. Bergen, M. Lange, S. Peidli, F. A. Wolf, and F. J. Theis, “Generalizing RNA velocity to transient cell states through dynamical modeling,”Nat. Biotechnol., vol. 38, no. 12, pp. 1408–1414, 2020.

[52] Y. Zhang et al., “Influenza Research Database: An integrated bioinformatics resource for influenza virus research,”Nucleic Acids Res., vol. 45, no. D1, pp. D466–D474, 2017.

[53] I. Letunic and P. Bork, “Interactive Tree of Life (iTOL) v4: Recent updates and new developments,”Nucleic Acids Res., vol. 47, no. W1, pp. W256–W259, 2019.

[54] UniProt Consortium, “UniProt: A worldwide hub of protein knowledge,”Nucleic Acids Res., vol. 47, no. D1, pp. D506–D515, 2019.

[55] Y. Shu and J. McCauley, “GISAID: Global initiative on sharing all influenza data – from vision to reality,” Eurosurveillance, vol. 22, no. 13. p. 30494, 2017.

[56] S. Guindon, J. F. Dufayard, V. Lefort, M. Anisimova, W. Hordijk, and O. Gascuel, “New algorithms and methods to estimate maximum-likelihood phylogenies: Assessing the performance of PhyML 3.0,”Syst. Biol., vol. 59, no. 3, pp. 307–321, 2010.

